# Bending friction: a new mechanism of dissipation within DNA explains its slow looping dynamics

**DOI:** 10.1101/2025.07.10.664086

**Authors:** Georgii Pobegalov, Maxim Molodtsov, Bhavin S. Khatri

**Affiliations:** The Francis Crick Institute, London, United Kingdom; Dept of Physics & Astronomy, University College London, United Kingdom; Department of Life Sciences, Imperial College London, United Kingdom

## Abstract

DNA bending and looping is crucial for gene expression, packaging, and chromatin organisation, as well as the design of artificial nanomaterials and devices. But what determines how quickly DNA bends? While DNA’s static flexibility is well-characterised by its persistence length, we lack an understanding of how quickly DNA responds to mechanical forces: remarkably current semiflexible polymer theory based on solvent dissipation underestimates spontaneous looping times by ~1000-fold. By analysing fluctuations of DNA several kilobases long and developing new theory for bending dissipation in semiflexible polymers, we show DNA bending dynamics cannot be explained by solvent friction alone and requires significant contributions from intramolecular friction. The theory defines a new material constant of DNA — the bending friction, which we determine to be *ζ_B_* = 241 ± 17 *μ*g nm^3^/ms. Strikingly, our measurement does not depend on the buffer ionic conditions. We predict bending friction will dominate DNA dynamics between ≈ 50 nm and 420 nm and significantly longer under external force. We show that mean first passage time calculations are greatly simplified when bending friction dominates and so using this constant, with no fitting parameters, we accurately predict the slow experimental spontaneous looping times. Our discovery of significant bending dissipation is unexpected as DNA has no obvious large (> *k*_*B*_*T)* internal energy barriers. The salt-independence of this dissipation also rules out long range electrostatic interactions as its origins. Instead our findings point to a complex local energy landscape for bending and a potential previously unappreciated role of water binding DNA constraining its local mobility. Our findings radically change our understanding of DNA dynamics and reveal DNA as a viscoelastic semiflexible polymer with dramatically slower dynamics compared to an ideal elastic rod. This work establishes bending friction as a fundamental material property that must underpin any model of DNA dynamics in biology, physics, and nanotechnology.

## I. INTRODUCTION

DNA is one of biology’s most important polymers, storing the instructions for life. Beyond information storage, its mechanical properties play a fundamental role in numerous processes in molecular biology, from transcription initiation [7, 41] and chromatin packing around nucleosomes [45], to genome organisation in eukaryotes [24, 37, 50]. These processes often involve molecular motors that manipulate, bend and loop DNA, where there is evidence that the local bending properties of DNA may be coded through sequence in the genome[3]. The spontaneous bending and looping of DNA is also a crucial step in high throughout sequencing through the process of bridge amplification [5], which forms the basis of the recent advances in genomics and its industry and the emergence of DNA as a means high volume data storage [2]. In addition, our ability to artificially construct DNA nanostructures and develop bio-inspired nanomachines [21, 25, 47] depends crucially on understanding its mechanical properties.

While DNA’s bending elasticity determines its equilibrium shape under mechanical constraints or forces, we lack a comparable understanding of an equally important mechanical property: how rapidly DNA changes shape under external forces or thermal fluctuations. The prevailing framework for DNA dynamics, semiflexible polymer theory — that assumes dissipation that limits dynamics arises solely from Stokes friction with the solvent[27, 36] – severely underestimates spontaneous loop-closure times for short DNA (~100nm contour length or 300bp) by a remarkable 1000-fold [32, 51]. Therefore, we lack a theoretical framework to explain these slow looping dynamics and so are unable to answer the simple question: what determines how quickly DNA bends.

However, Kuhn’s theorem has long predicted that for short flexible chains, intramolecular friction – arising from internal energy barriers — must dominate polymer dissipation[9, 13, 28, 31, 38, 39]. de Gennes illustrated this with a striking thought experiment: if thermal energy *k*_*B*_*T* is much smaller than internal energy barriers Δ*U*^‡^, a single polymer chain would enter a frozen, glassy state[13]. Lacking the obvious internal rotation barriers of flexible polymers, DNA is often treated as a simple elastic rod. Yet, its complex molecular structure creates a potentially intricate energy landscape for bending, shaped by atomic interactions within DNA and with surrounding water (Fig.1). Whether these processes within DNA are sufficient to explain empirically measured loop-closure times is unknown.

**FIG. 1.**
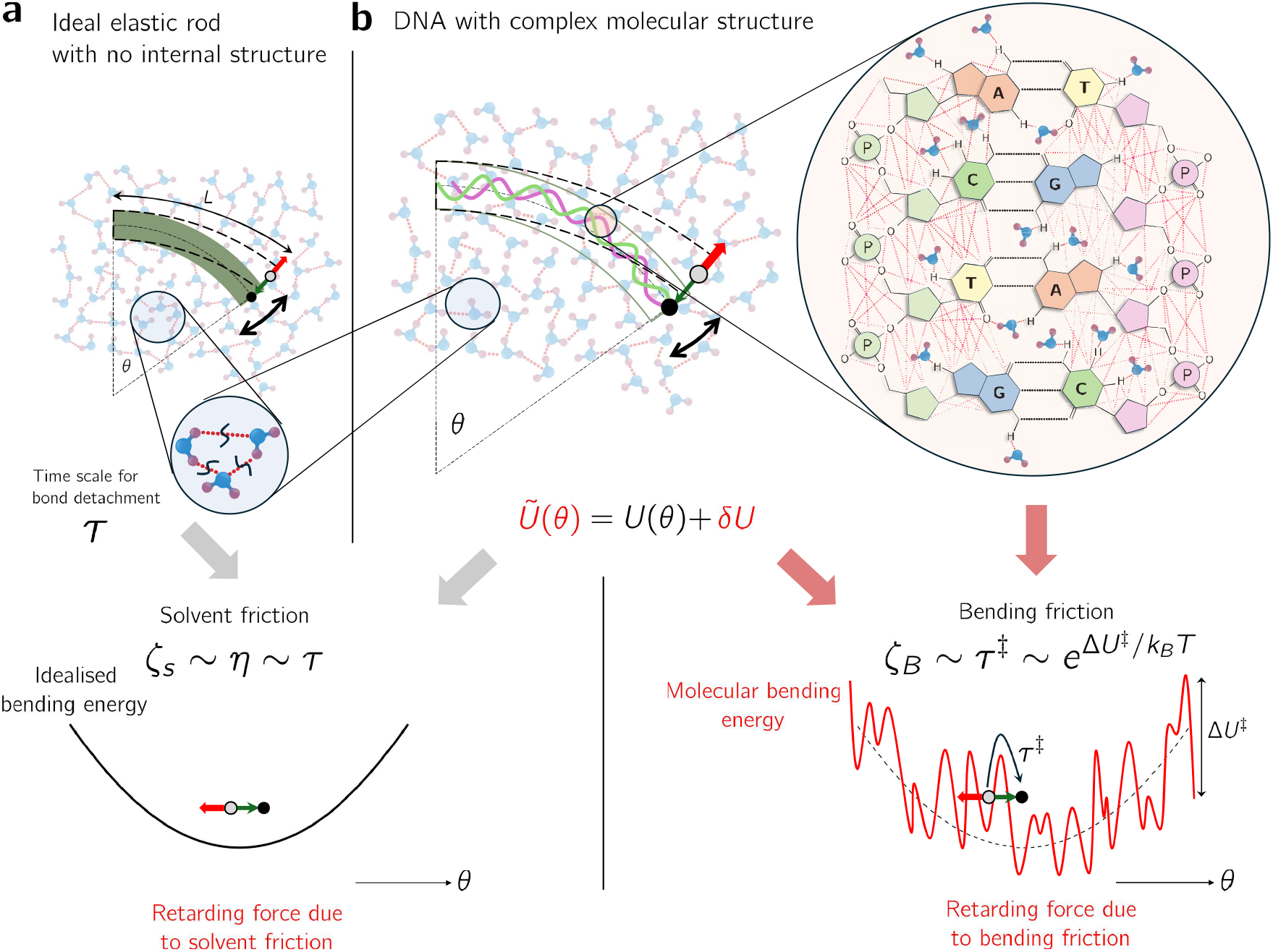
The origins of bending friction within DNA. a) an ideal elastic rod with no internal structure has a smooth quadratic elastic energy *U* (*θ*) as function of bend angle ***θ***, where the curvature is controlled by the bending elasticity. Rod friction *ζ*_*s*_ arises from viscosity with solvent, determined by the timescale *τ* of water hydrogen bond lifetimes. Grey dot moving to black dot over time indicates a change in bend angle with red arrow indicating retarding frictional force. b) DNA segment: unlike an idealised elastic rod has a complex internal molecular structure. As DNA bends, complex changes in the relative positions of all the atoms, gives a corresponding complex change in the energy of DNA due to numerous interactions (red dotted lines). In addition, specific binding of water molecules to nucleotides, will necessitate unbinding and binding of water as DNA bends. These factors could give rise to rough energy landscape (red) with local energy barriers, where Δ*U*^‡^*/k*_*B*_*T* is the roughness of the energy landscape – on the scale of thermal energy *k*_*B*_*T* – to DNA bending due to local interactions in DNA, and *ζ*_*B*_ is the emergent bending friction of DNA due to local barrier hopping on timescale *τ*^‡^.

By analysing DNA fluctuations across a range of lengths and tensions, and developing a new first-principles theory for bending dissipation in semiflexible polymers under tension, we reveal clear signatures of intramolecular friction that rule out a purely solvent-based origin. What emerges is a new material constant of DNA – the bending friction – describing the local dissipation within DNA itself as it bends. Strikingly, we find this constant is independent of the buffer’s ionic strength, which suggests its origin lies in local energetic interactions within DNA and potentially water binding DNA, which would be a new unappreciated role for water modulating its bending dynamics. Using our measurement of this constant, we show that we can fit measured loop-closure times with no fitting parameters. This demonstrates that DNA’s slow looping dynamics is explained by intramolecular bending friction and radically changes our picture of DNA at these length scales: from an ideal elastic rod to a viscoelastic semiflexible polymer with dramatically slower dynamics.

## II. EXTRACTING DNA FRICTION FROM SINGLE-MOLECULE FLUCTUATIONS

To determine the role of bending friction in DNA dynamics, single DNA molecules were tethered between two beads each in optical traps and stretched to various mean forces *F* (1pN≤ *F* ≤ 5pN) by appropriately choosing the distance between the trap centres Fig.2a. This range ensures DNA is highly stretched *(F* ≫ *k*_*B*_*T* / ℓ_*P*_ ≈ 0.08 pN) while avoiding backbone extension. At each force, we recorded the positions of both beads, *x*_1_(*t*) and *x*_2_ (*t*), at 78 kHz for 6 seconds. DNA length fluctuations are calculated as ℓ(*t*) = *x*_2_(*t*) − *x*_1_(*t*) + *X* − *D*, where *X* is the separation between trap centres and *D* the diameter of the beads. Example time series of ℓ(*t*) are shown in Fig.2b as a function of increasing force. To specifically isolate dissipative dynamics, we calculated the velocity power spectral density (VPSD, *P*_*ν*_(*ω*)), derived from the length fluctuations as *P*_*ν*_ = *ω*^2^ *P*_ℓ_ (Fig.2c), where *P*_ℓ_(*ω*) is the regular power spectrum. This transformation weights higher frequencies, providing a direct measure of the power dissipated unit frequency – which increases at higher frequencies – and thus a more reliable signature of DNA friction (Fig.S1).

**FIG. 2.**
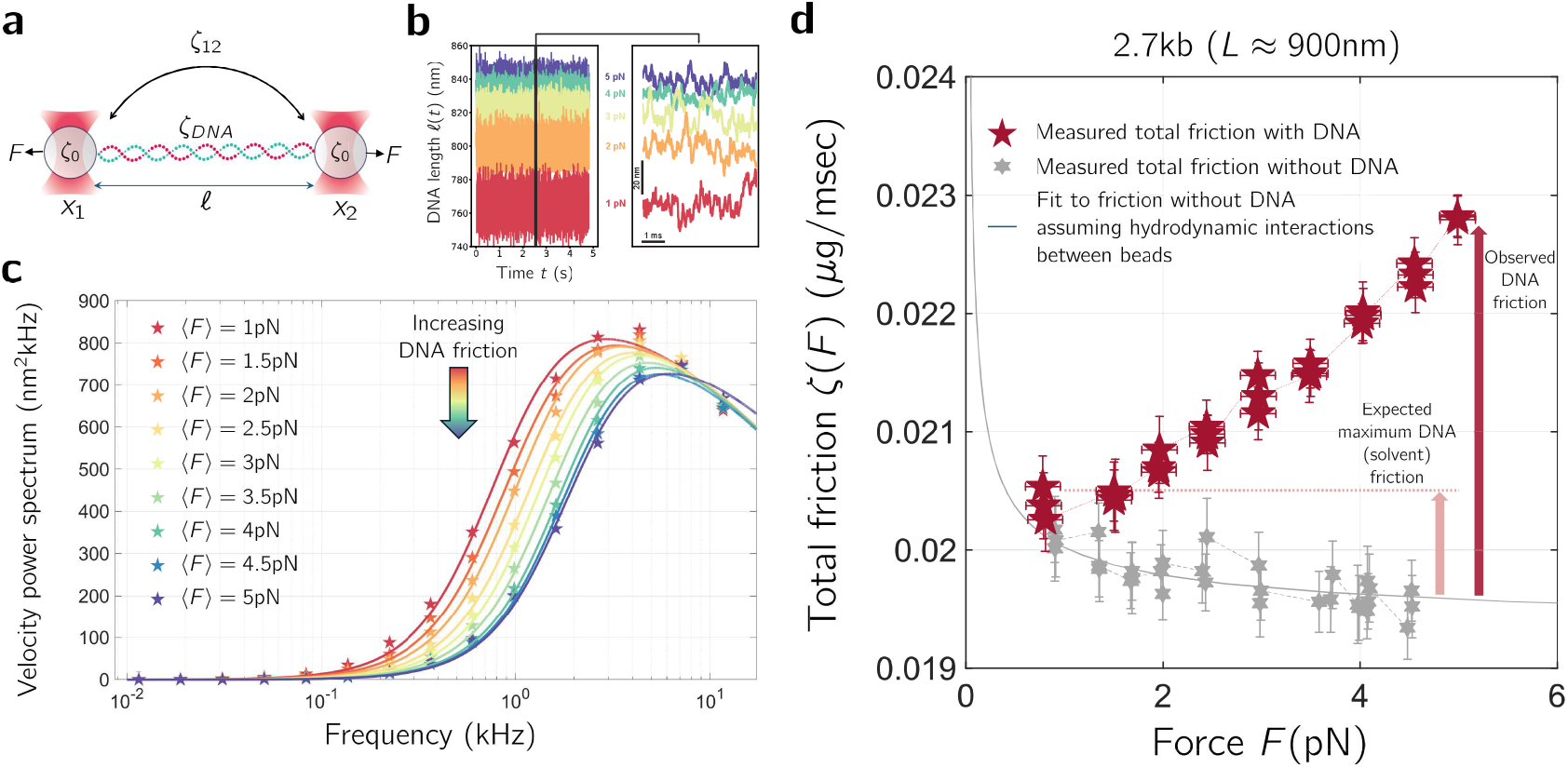
Measuring the friction response of DNA as a function of force. a) DNA is tethered between two beads at a mean force F using dual optical trap. Time series of the position of each trap, *x*_1_(*t*) and *x*_2_(*t*), is recorded, where total friction of system is given by 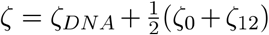 (Supplementary Information:Eqn.Sl7), where *ζ_DNA_* is the friction of DNA, *ζ*_0_ friction of each bead and *ζ*_12_ the friction due to the hydrodynamic interactions between the beads. b) Example time series of length ℓ (*t*) for 2.7kbp (903 nm) DNA as a function of force, showing increased extension for increasing forces. c) the velocity power spectrum (VPSD) of the fluctuations in b): decrease in peak height with increasing tension can only be explained by increasing friction of DNA (Supplementary Information: section S3). d) Total effective friction of the trap DNA system (red pentagrams) as a function of force. DNA friction increases above background hydrodynamic friction (grey hexagrams – measured without DNA – and grey solid line, fit with hydrodynamic theory between two beads). This contrasts to expectation if solvent friction were only source of dissipation (pink dashed line: 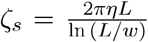 [16]; *L* = 902.6 nm, *w* = 2.4 nm[34]).

The velocity power spectrum is a function of two main force-dependent vis-coelastic parameters of DNA: the end-to-end elasticity *K*_*DNA*_*(F)* – which is the well-characterised entropic elasticity often described by a Worm-like chain (WLC) model [8, 35] – and the end-to-end friction of DNA *ζ*_*DNA*_(*F*). Our data in Fig.2c shows that the velocity power spectrum is highly sensitive to changes in the aver age tension applied to DNA. As we demonstrate in Fig.S2 the maximum value of the VPSD is a direct read-out of the inverse of the total friction constant of the system. Therefore, even before fitting any model to the VPSD, we can see that the friction of DNA is increasing with increasing tension, which as we argue below cannot be explained by solvent friction. Fitting to the VPSD at each force in Fig.2c, using a model of fluctuations which includes appropriates high frequency corrections to Brownian motion (Supplementary Information: Eqn.S18), we determine the combined friction constant of the bead, trap and DNA, *ζ* (*F*), as a function of force shown in Fig.2d, as solid squares.

Our data show that the friction constant with DNA tethered between beads increases approximately linearly with increasing force over and above the back-ground bead friction (grey hexagrams with solid grey line fit to Oseen hydrodynamic theory [17] (Supplementary Information Eqn.S11)). This increase cannot be attributed to the solvent friction of DNA: at high stretch *(F* ≫ *k*_*B*_*T* / ℓ_*P*_ 0.08pN) solvent dissipation gives an approximately constant friction of *ζ*_*s*_ 0.00092*µ*g/ms for 2.7kb long DNA (dashed pink line in Fig.2d, where we assume DNA is a rod of length *L* ≈ 900nm, and width *w* = 2.4 nm[34]). Thus, contrary to the predictions based on the solvent friction alone, our data in Fig.2d show that friction increases with force and reaches approximately 0.0035µg/ms, above the background hydrodynamic friction, at *F* = 5 pN, which is almost four-times the expected solvent friction. In the next section, we develop a theory of bending dissipation in DNA to explain these observations.

Our fitting also yields the elastic constant *κ*(F), which matches the entropic elasticity expected of a WLC at high stretch (Fig.S3). The fit of the VPSD data also accounts for the effective frequency-dependent mass of water entrained around each bead, which should be independent of force, which we verify in Fig.S4.

## III. FORCE-DEPENDENT FRICTION IS EXPLAINED BY BENDING DISSIPATION WITHIN DNA

Because solvent friction alone could not explain the linear increase in friction of DNA with force, we sought to rationalise our findings by modifying the standard model used to describe DNA dynamics when highly stretched at a tension *F (F* ≫ *k*_*B*_*T* / ℓ_*P*_ 0.08pN). Semiflexible polymer dynamics describes DNA as an idealised elastic rod with space curve ***R(s***, *t)* – wheres is the backbone co-ordinate of the polymer — whose shape fluctuates dynamically due to Brownian motion and friction with the solvent. We propose that due to internal frictional processes within DNA there will be local bending dissipation proportional to the rate of change of local curvature, described by the Rayleigh dissipation function 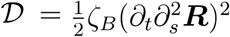, where *ζ*_*B*_, is the bending friction constant[23, 43]. Calculating the local forces due to this dissipation gives rise to a modified equation for semiflexible polymer dynamics under tension, which we call the Dissipative Worm-Like Chain (DWLC):

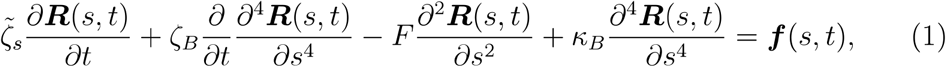

where 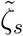 is a solvent friction per unit length, and *κ*_*B*_ = *k*_*B*_*T*ℓ_*p*_ is the bending elastic constant and ***f(s***,*t)* is a temporally white, but spatially coloured, noise term as is necessary for a continuum description of internal friction[29] (see Supplementary Information section S2 for details).

Our aim is to evaluate the additional, or excess, end-to-end friction of the chain, as a function of tension *F*, due to bending dissipation, over and above hydrodynamic solvent contributions or Stokes’ friction of DNA. We do this by analysing Eqn.1 using standard normal mode analysis to give the mode relaxation time for the *n*^*th*^ mode:

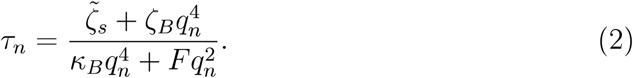

where *q*_*n*_ is its wavenumber. We then evaluate the relevant *4*^*th*^ order moments using Wick’s theorem[l 7] to calculate the autocorrelation function of the end-to end distance Δ*R* of the chain:

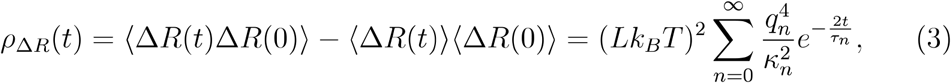

where mode elasticity is given by 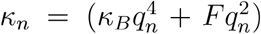 and mode friction by 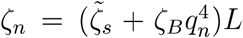. The end-to-end excess friction is then calculated from the derivative of the autocorrelation function [18]:

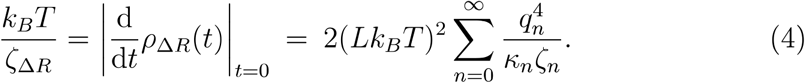

We then take the continuum limit with *q*_*n*_ → *q*, with mode spacing *δ*_*q*_ = *π* / *L* to convert the summation to an integral to give the following, valid for forces *F* > 0.08pN, in the limit of the chain at high force/stretch (full derivation given in Supplementary Information section S2):

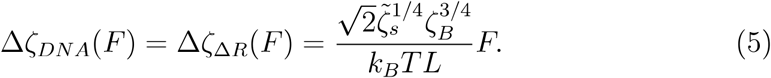

This shows that explicit inclusion of bending friction in the equations of motion leads to the observed linear increase of the friction coefficient with external force (Fig.2d). Note that the friction of the dissipative WLC is inversely proportional to the contour length *L*, which is a classic signature of internal dissipation in polymers [13, 29]. Thus, DNA friction that takes into account internal bending dissipation increases with force, and it does so more significantly for shorter DNA.

## IV. A NEW MATERIAL CONSTANT OF DNA

Having established the theoretical description of bending dissipation using the DWLC model, and parameterised by the bending friction constant, we can now infer its value from the experimental data. Since bending dissipation in DNA gives rise to a friction inversely proportional to contour length, we measured a number of single molecules of DNA for lengths between 2.7kbp and 8.8kbp (≈900nm to 3000nm). To extract the DNA friction from the measurements, we determined the excess DNA friction, Δ*ζ*_*DNA*_(*F*), as the residual friction above the background friction, due to the hydrodynamic interactions between beads (grey pentagrams in Fig.2d) and the hydrodynamic (Stokes) solvent friction of DNA with a fitting parameter to account for the complex hydrodynamic interactions between the beads and DNA (Fig.S6). The data is then fit using Eqn.5, which has a single fitting parameter *ζ*_*B*_.

The resulting measurements of the excess friction for all DNA lengths (various colours), with the effective background bead and DNA solvent friction removed are shown in Fig.3. The fitting of Eqn.5 to the experimental data shows that our DWLC model accounts for the main unexpected feature of the data: a linear increase of the excess DNA friction with increasing force. In addition, Fig.3 shows that as the contour length decreases, the excess friction increases, which is in qualitative agreement with Eqn.5. In Fig.S7, we additionally plot the normalised friction Δ*ζ*_*DNA*_ as a function of force, which shows excellent quantitative agreement with 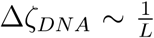. This is in clear contrast to solvent DNA friction, which would increase approximately linearly with increasing length (Fig.S5). Thus, the experimentally observed excess friction is fully explained by the bending friction of DNA.

**FIG. 3.**
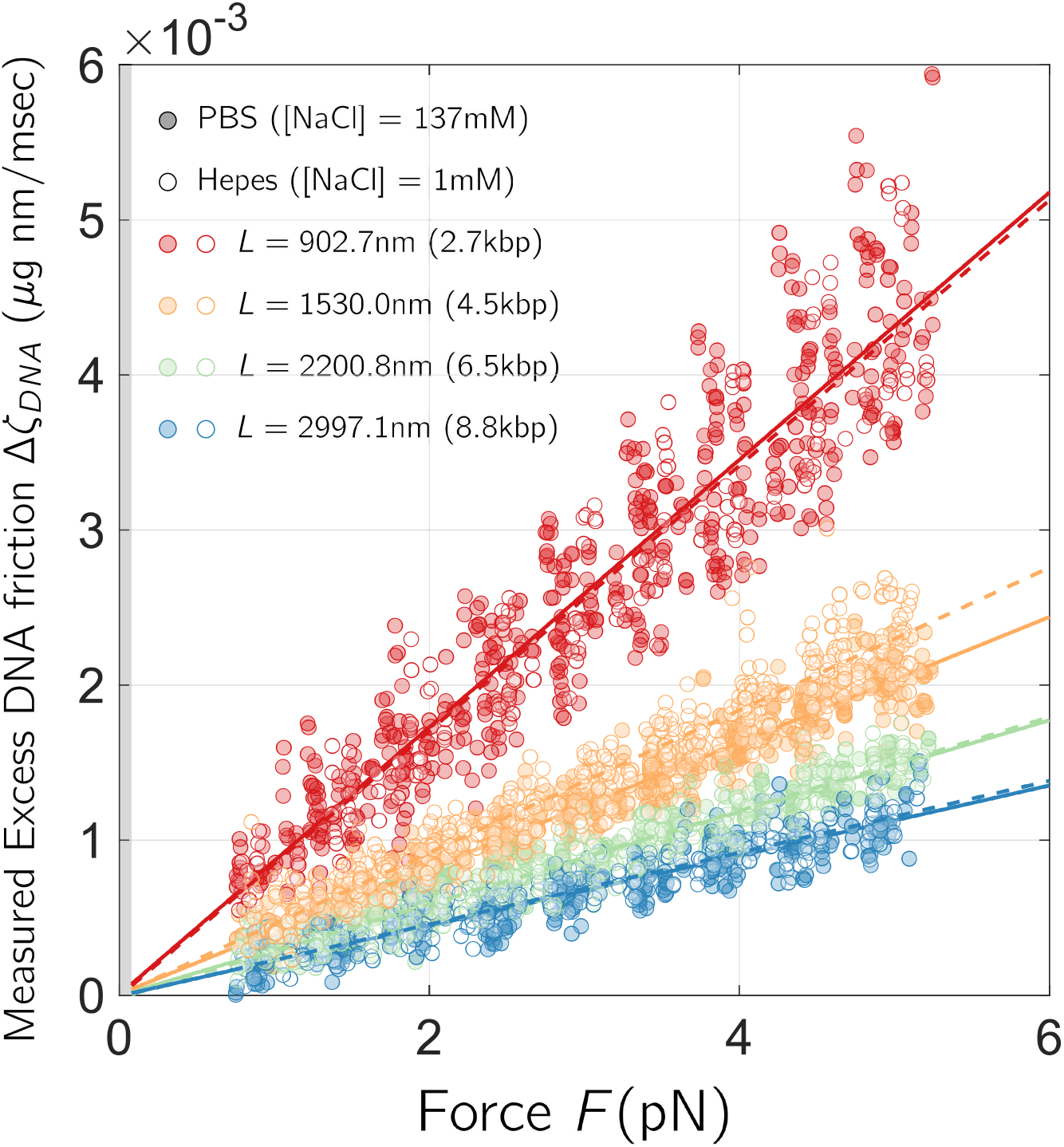
The end-to-end friction of DNA. The measured excess end-to-end friction of DNA as a function of force showing linearity and increasing friction for decreasing contour length as predicted by Eqn.5. Closed circles are in normal PBS buffer and open circles are in the low salt Hepes buffer. The data is fit using Eqn.5 with single fitting parameter *ζ*_*B*_, with 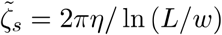 with fits shown with solid lines (PBS) or dashed lines (Hepes). The number of measurements on independent single molecules at each length and for each buffer condition are given in Methods (Table.II). The shaded region indicate the forces for which we would expect the linear dependence to be invalid (i.e. when at low stretch when *F* ≪ *k*_*B*_*T/*ℓ_*P*_ ≈ 0.08 pN); in fact as *F* → 0, we would expect the excess friction due to bending dissipation to tend to a constant. These measurements cannot be due to solvent dissipation, as we would expect solvent friction to be approximately independent of force at high stretch and increase with increasing contour length of DNA (Fig.S5).

Furthermore, to test whether DNA bending friction depends on long-range electrostatic interactions within DNA, we compared experiments performed at physiological salt concentrations (in PBS buffer) with experiments carried out under very low salt conditions (1 mM NaCl, in 1mM Hepes buffer; see Methods). This comparison (closed and open data circles, respectively, in Fig.3) shows no dependence of the bending friction on the salt concentration (Fig.S7). As a control, for each molecule in these experiments, we also performed force-extension measurements and extracted their persistence lengths. As expected and in quantitative agreement with previous studies[4, 10, 22, 52], we observed an increase in the persistence length for the low salt buffer (Supplementary Information: Fig.S12).

Our experiments show that bending friction is a material constant of DNA independent of the buffer conditions. This suggests that the origin of the excess DNA friction we measure is due to local interactions within DNA, rather than long-range electrostatic interactions. Therefore, we combine fits to data across both buffer conditions, accounting for finite-size effects [48] (Fig.S7), to give an empirical value of *ζ*_*B*_ = 241 ± 17*µ*g nm^3^/ms (s.em).

## V. THE DYNAMICS OF DNA LOOPING ON SMALL LENGTH SCALES

We have determined a new fundamental material constant for DNA, *ζ*_*B*_ which determines how DNA locally dissipates energy as it bends and loops. With our measurement of *ζ*_*B*_ we can now quantify the relative contributions of solvent and bending friction for any DNA length at zero force. The normal mode analysis (Eqn.2) shows that the critical length scale at which solvent and bending contributions to the friction are equal is *L*^*^ ≈ 750nm (Fig.4a) (Supplementary Information: section S4), which is equivalent to a DNA length of approximately 2.2kbp. This figure also shows that for DNA contour lengths *L* ≤ 420nm, solvent friction is 10 × smaller than bending friction; in other words we can model the spontaneous dynamics of DNA, considering only the contributions from bending friction, for contour lengths up to about *9*ℓ_*p*_, or 1.2kbp. For very small contour lengths less than persistence length, DNA kinking likely plays a significant role in DNA flexibility and dynamics[53]. Therefore, we expect bending friction dominates for DNA lengths ℓ_*P*_ ≤ *L* ≤ *9*ℓ_*P*_ and have significant contributions up to 15ℓ_*P*_ → 20ℓ_*P*_ (750nm → l*µ*m, or 2.2kbp → 3kbp) at zero force. As we show below, the stochas tic dynamics of DNA is greatly simplified in the regime where bending friction dominates.

**FIG. 4.**
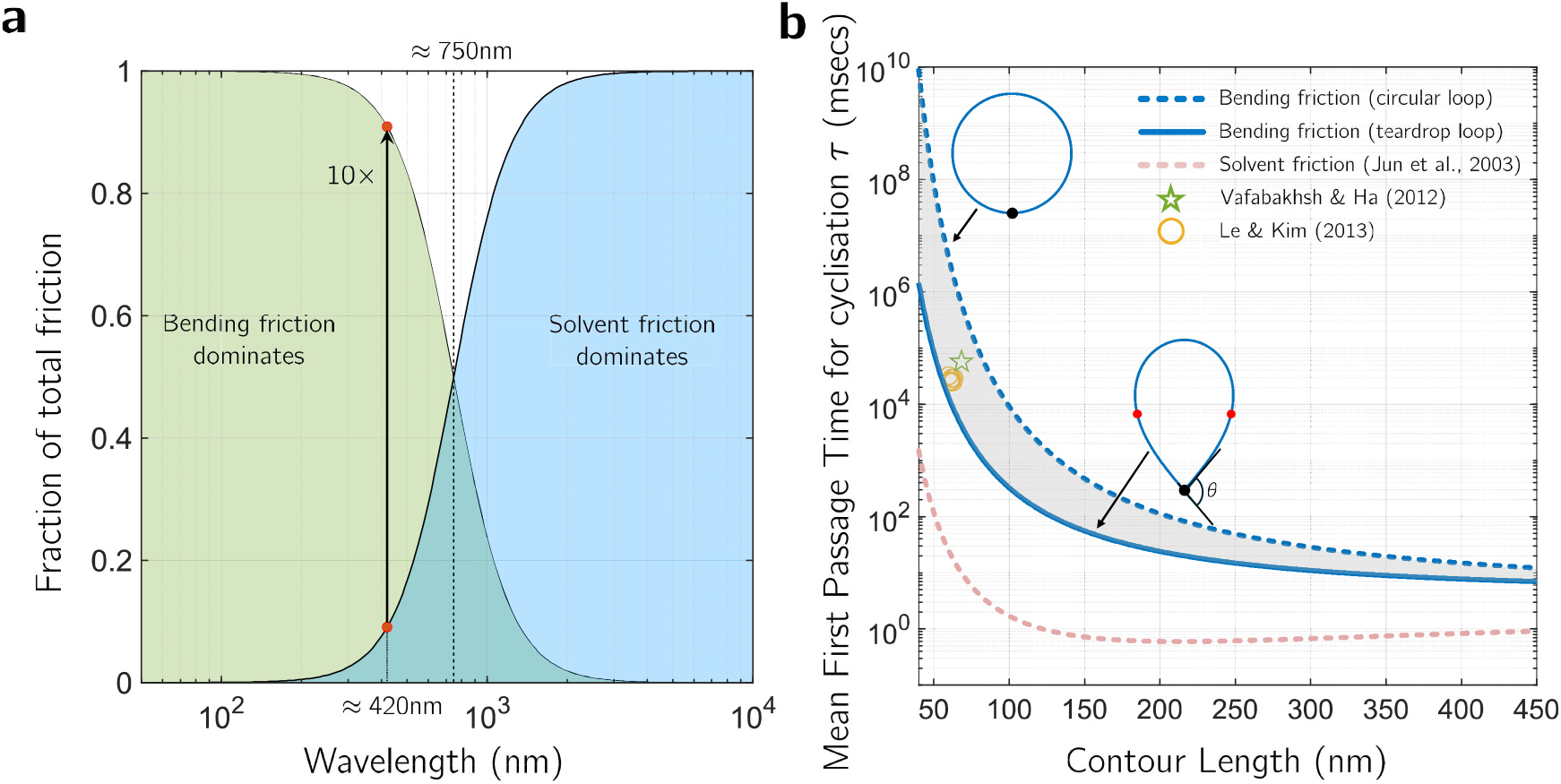
DNA dynamics on small length scales. a) Relative contribution of bending and solvent friction to total mode friction for a given wavelength. b) Plot of mean first passage times (MFPT) for DNA loop closure as a function of contour length. The predictions based on the theory developed in this paper for circular loops (Eqn.S3) and minimum energy tear-drop loops (Supplementary Information: section S5), where bending friction and bending energy dominate, are shown as dashed blue line and solid blue line, respectively. While the prediction based on a WLC with only solvent dissipation [27] is shown by the dashed pink line. Experimental measurements of MFPT using cyclisation assays are shown by open circles[32] and open pentagrams [51].

From this zero force limit, as tension increases and the entropy of the chain decreases, we expect in general that the bending friction contribution will increase, and therefore be more important at even longer length scales. Eqn.5, demonstrates this explicitly once the chain is at high stretch *(F* ≫ *k*_*B*_*T* / ℓ_*P*_ ≈ 0.08 pN). In cell nuclei DNA can be locally subjected to forces up to 20 pN [42, 44], at which we calculate bending friction will dominate up to ≈ 1.2*µ*m and with significant contributions up to ≈ 12*µ*m (Fig.S8). At lower forces and stretch, we expect the bending friction contribution to decrease monotonically to the zero force limit described by the mode analysis above.

When bending friction dominates the dynamics, we can calculate a new useful quantity – the persistence time of DNA – which is the average time it takes for DNA to completely change shape due to random thermal fluctuations. In the absence of external force and in the limit that bending friction dominates, this is simply given by the mode relaxation time, since all modes relax with the same characteristic time *τ* _*n*_ independent of mode number *n*:

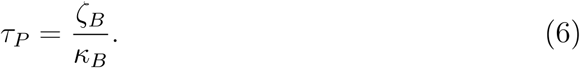

Given the value of *ζ*_*B*_ that we have determined, and assuming *κ*_*B*_ = *k*_*B*_*T* / ℓ_*P*_ ≈ 200pNnm^2^ this gives *τ*_*P*_ = l.2ms. Thus, DNA segments of lengths 750nm (1.2kbp) or shorter completely change their shape under thermal fluctuations in approximately 1.2 ms. Furthermore, as we will see below, the persistence time of DNA is directly proportional to the time it takes DNA to spontaneously make a loop.

### A. Comparison to experimentally measured DNA looping times

Cyclisation assays have become the standard way to estimate the rate of loop closure of DNA [32, 51], the characterisation of which is important to understanding the dynamics of number of DNA looping processes in molecular biology. Current theory to calculate mean first passage times are based on semiflexible polymer dynamics without bending friction [27, 36] and are consequentially complex calculations as they require consideration of the different time scales of each of the modes contributing to the polymer dynamics. However, when bending friction dominates solvent friction and in the limit that bending energy dominates the chain entropy, we can make the assumption that the DNA conformation is roughly an arc of a circle with curvature Γ = 1/ *R*, where *R* is the radius of this circle. The resulting stochastic dynamics is then given by the standard Smoluchowski equation for a single degree of freedom, the curvature Γ, with effective friction *ζ*_Γ_ = *Lζ*_*B*_ and bending energy 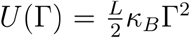 Using the standard flux-over population method and Kramer’s approximations (see Supplementary Information– section S5 – for details) we find the mean first passage time (MFPT) τ^*^ for loop closure is:

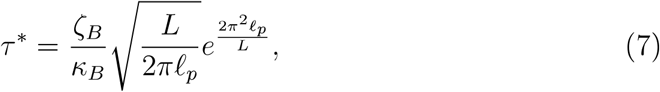

which is a pleasingly simple closed-form expression, compared to the theory of Jun et. al [27] for a WLC where only solvent dissipation is considered. The ratio of bending friction to bending elasticity is simply the persistence time of DNA (Eqn.6), which means the loop closure time is proportional to the persistence time of DNA conformations.

However, experiments measuring the time for loop closure of DNA [26], suggest that it is most likely to adopt – at least initially – a tear-drop conformation, where the ends come together at a non-zero angle *θ* between the end tangent vectors (Fig.4b). Previous theoretical work, shows the minimum energy conformation corresponds to *θ* ≈ 100° [54] and an overall energy which is roughly 70% the energy of a circular loop [1, 54]. In Supplementary Information (section S5), we estimate the time for closure of a tear-drop loop, by solving the Euler-Lagrange equations for a rod with free ends to calculate its curvature as a function of backbone coordinate. We then make the assumption that the MFPT will be dominated by a portion of the chain around its midpoint where the curvature is highest, which are indicated by the red circles in Fig.4b.

In Fig.4b, we compare the circular and tear-drop predictions of loop-closure times – based on bending dissipation with bending friction constant we measured experimentally – against theory assuming only solvent friction (Jun et al. [27]) and experimentally available looping times from Vafabakhsh & Ha [51] and Le & Kim [32], for DNA lengths between 60 − 70nm. Our theory agrees with the data, which lies between the lower bound set by the tear-drop loop closure theory and upper bound by the circular loop closure theory; in contrast, we see that the theory with only solvent friction massively underestimates the looping time by approximately 1,000-fold. This is a remarkable agreement between experimental looping times and the prediction of bending dissipation within DNA given no fitting parameters.

## VI. DISCUSSION

A key question remains: what is the underlying origin of bending friction within DNA? Our finding that the bending friction constant is insensitive to salt concen-tration, and the strength of electrostatic screening, suggests long-range electro-statics, such as repulsion from the negatively charged backbone, plays a weak role. We instead attribute the origin to local complex energy barriers from the complex atomic interactions within each base pair (Fig.1) as well as potentially complex interactions with water. The latter contributions may come from water binding in the minor groove [30, 46] or specific hydrogen bonds, such as adenine-thymine bridges predicted to stabilize DNA by ≈3 *k*_*B*_*T* [33]. This previously unanticipated role of water in bending friction would carry important implications for the proper choice of water models to obtain realistic explicit solvent molecular dynamics simulations of DNA, as well as implicit solvent and coarse-grained simulations of DNA, such as oxDNA[40].

Our work identifies a new material constant for local bending dissipation within DNA which we have demonstrated dominates looping and bending dynamics from 50nm to 420nm (150 bp to 1.2 kbp), with significant contributions up to ≈ 1*µ*m (3 kbp) and potentially an order of magnitude longer under physiological tensions of~ 20pN found in the cell [42, 44].This has broad implications for the trade-off between the maximum speed versus energy demands of biological processes where proteins, enzymes, or motors manipulate, bend or loop DNA. Examples at these length scales in biology include gene expression[ll, 19, 20] and loop initiation and extrusion in replisomes, SMCs or telomerase complexes[12, 14]. Our calculations in Fig.4b show that for DNA between 50nm and 420nm loop-closure times vary by many orders of magnitude, ranging from roughly 3 hours to 10 ms. This suggests spontaneous loop formation for small length DNAs is likely inefficient and therefore requires significant energy input to overcome bending dissipation, whereas for longer DNA, looping may occur spontaneously. These same trade-offs will play a role in the design, construction and optimisation of nanomaterials and devices[47] – for example strand-displacement artificial DNA motors[49, 55] – and particularly in the engineering of bio-inspired systems that mimic the non-equilibrium nature of molecular biology[21], where dynamics will be a key consideration.

Overall our findings show that DNA – despite lacking any obvious large internal energy barriers – nonetheless has significant intramolecular dissipative forces described by a new physical principle of bending dissipation in semiflexible polymers, which explain the puzzle of its very slow loop-closure times. This sets-forth a new paradigm for DNA dynamics – the dissipative worm-like chain (DWLC) – and calls for investigation of signatures of bending dissipation in other semiflexible polymers such as actin and microtubules. This work establishes bending friction as a fundamental material property that must underpin any model of DNA dynamics in molecular biology, DNA-based condensates[15] and in material science for its implications for defining new engineering principles of DNA nanodevices and nanomaterials[21, 25, 47].

## Supporting information

Supplementary figures and derivations

## ACKNOWLEDGEMENTS

We would like to thank Ard Louis, Agnes Noy, Tanniemola Liverpool and Frank Uhlmann, for various discussions and suggestions on the text and figures of the manuscript. We thank Holly Folkard-Tapp for her advice on the design of the graphical elements.

## AUTHOR CONTRIBUTIONS

GP performed the experiments. MM contributed to study design, supervised and provided funding for the experiments. BSK conceived and designed the study, and performed the theoretical calculations. BSK and MM wrote the manuscript with input from GP.

## Appendix A: Methods

### 1. DNA functionalisation

To facilitate attachment to beads, shorter DNA molecules (2. 7 kbp and 4.5 kbp) were labelled with a single biotin and a single digoxigenin at either end of the DNA molecule. For longer molecules (6.5 kbp and 8.8 kbp) both DNA ends were labelled with biotins. Restriction enzymes, DNA polymerases and buffers were purchased from New England Biolabs (NEB). Oligonucleotides were purchased from Merck. All PCR reactions were performed using Phusion High-Fidelity DNA Polymerase in a Phusion HF Buffer (NEB).

#### a. 2,655 bp biotin-digoxigenin DNA

2,655 bp DNA fragment was synthesised by PCR using Lambda-phage DNA (NEB) as a template, a 5’-biotin containing primer (LHD2_RT7bio) and a primer containing Xbal restriction site (LHD2_FXbaI). PCR reaction was purified using a PCR Cleanup Kit (Monarch T1030S, NEB) and the product was further digested with Xbal restriction enzyme (NEB) in a Standard Taq Reaction Buffer (NEB) for 40 minutes at 37 °C to generate a 4-nt single-stranded overhang. Xbal was inactivated by incubating the reaction for 20 minutes at 65 °C and the second end of the DNA was labelled with digoxigenin by end filling reaction using Taq DNA polymerase (NEB) and a mixture of dATP, dCTP, dGTP (Promega) and dUTP-digoxigenin (Jena Bioscience) performed for 30 min at 72 °C. The final product was purified using Micro Bio-Spin P30 spin column (Bio-Rad).

#### b. 4,500 bp biotin-digoxigenin DNA

4,500 bp DNA fragment was synthesised by PCR using Lambda-phage DNA (NEB) as a template, a 5^′^-digoxigenin containing primer (LHT1_Fdig) and a 5^′^ biotin containing primer (4.5kb_rev _bio). PCR reaction was purified using a PCR Cleanup Kit (Monarch T1030S, NEB).

#### c. 6,473 bp biotin-biotin DNA

9,847 bp plasmid DNA (8×601-pKYB1, a gift from Graeme King) was double digested with XbaI and Spel restriction enzymes (NEB) for 1 hour at 37 °C in rCutSmart buffer (NEB), followed by inactivation for 20 minutes at 80 °C. This resulted in 2 fragments (6,473 bp and 3,374 bp long) containing 4-nt single-stranded ends. DNA ends were further biotinylated by addition of Klenow Fragment (3 ^′^ →5^′^ exo-) of DNA Pol I (NEB), a mixture of dATP, dTTP, dGTP (Promega) and dCTP-biotin (Jena Bioscience) and incubation for 25 minutes at 37 °C, followed by inactivation for 20 minutes at 75 °C. DNA was cleaned-up from unincorporated nucleotides using Micro Bio-Spin P30 spin column (Bio-Rad). After tethering DNA to the beads only 6,473 bp-long molecules were selected by force-extension analysis.

#### d. 8,815 bp biotin-biotin DNA

8,815 bp plasmid DNA (2×601-pKYB1, a gift from Graeme King) was digested with XbaI (NEB) in the Standard Taq Reaction Buffer (NEB) for 45 minutes at 37 °C to generate a 4-nt single-stranded overhangs. DNA ends were biotinylated by end filling reaction using Taq DNA polymerase (NEB) and a mixture of dATP, dTTP, dGTP (Promega) and dCTP-biotin (Jena Bioscience) performed for 30 min at 72 °C. The final product was purified using the Micro Bio-Spin P30 spin column (Bio-Rad).

### 2. Single-molecule DNA tethering

Experiments were performed using commercial dual-trap optical tweezers (C-trap, Lumicks) combined with a multi-channel microfluidic laminar flow cell (u-Flux, Lumicks). Before experiments, the flow cell was passivated by incubating it with 0.5% Pluronic F-127 (Merck) diluted in PBS (Phosphate Buffered Saline) for at least 30 minutes. Pluronic was washed by flowing at least 1 mL of PBS through each channel of the flow cell. Individual DNA molecules were tethered between 2 polystyrene beads of 2 *µm* diameter. Shorter digoxigenin-biotin labelled DNA molecules (2.7 and 4.5 kbp) were first coupled to Anti-digoxigenin-coated beads (DIGP-20-2, Spherotech): 5 microliters of beads stock solution (0.1% w/v) were gently mixed with 5 microliters of DNA diluted down to 30 pM in PBS and incubated for 10 min. The mixture was further diluted by adding 300 microliters of PBS and loaded into the first channel of the flow cell. The second channel was loaded with PBS and third channel contained Streptavidin-coated beads (SVP-20-5, Spherotech) diluted down to 0.002% w/v in PBS. 4th and 5th channels of the flow-cell contained a low salt buffer (Hepes pH 7.5 1mM, NaCl lmM). To form a DNA tether, first a bead-DNA complex and a Streptavidin-coated bead were consecutively trapped in two optical traps in the first and the third channels respectively. The beads were further moved into the second channel where the free biotinylated end of DNA was attached to a Streptavidin-coated bead. The flow cell was extensively washed with PBS keeping only the 2nd channel open to remove any free-floating beads, after which the flow was stopped and measurements started. For longer biotinylated DNA molecules (6.5 and 8.8 kbp) the first channel contained Streptavidin-coated beads (SVP-20-5, Spherotech) diluted down to 0.002% w/v in PBS, second channel contained DNA diluted down to 5 pM in PBS, and the third channel was loaded with PBS. Two beads were trapped in the first channel and transferred into the second channel to form a DNA tether, which was further transferred into the 3rd channel and the flow was stopped prior to measurements. For measurements in the low salt buffer, after DNA fluctuations were measured in PBS, the 4th and 5th channels were extensively washed with the Hepes buffer (2 minutes at the pressure of 1 Bar). After this the flow was stopped and the DNA tether was transferred into the 5th channel where measurements were repeated. Not all DNA tethers lasted long enough to be measured in both PBS and Hepes buffers and hence in Table.II the number of single molecules for PBS and Hepes differ for each length. After molecule detachment, time series of bead positions were collected at approximately the same trap separations as the force measurements with DNA.

**TABLE 1.**
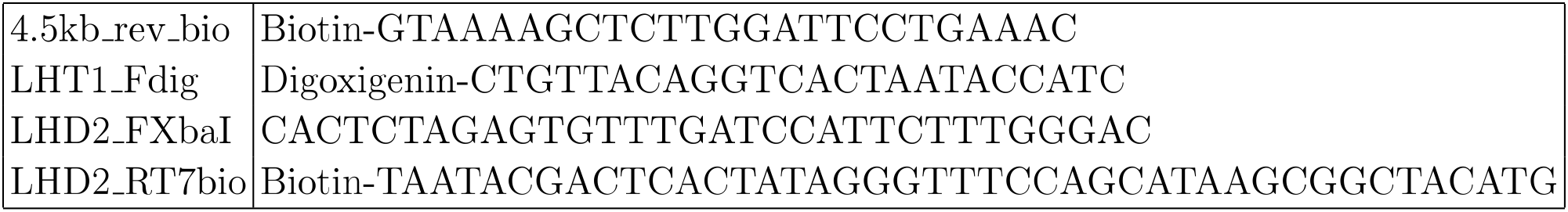
List of oligonucleotides.

**TABLE 2.** Number of single molecules used at each length and each buffer condition. The first number in the brackets are the number of single molecules used in PBS buffer and the second number the number used in the Hepes buffer. Each single molecule experiment consisted of measurements in PBS followed by Hepes; however, the Hepes stage did not complete for all single molecules and so the 2nd number is typically less than the first.

| Length | $n$ |
| --- | --- |
| 2.7kbp | (20,6) |
| 4.5kbp | (21,18) |
| 6.5kbp | (14,10) |
| 8.8kbp | (13,8) |

### 3. Force-extension and force-dependent DNA fluctuation measurements

After the DNA tether was formed and placed in a channel containing either PBS or a low salt buffer, a force-extension curve was recorded by increasing the beads separation at the rate of 40 nm per second while simultaneously recording the distance between the beads and the force in the optical traps until the DNA tension reached 10 pN.

After that the DNA molecule was fully relaxed by bringing the beads closer to each other. Subsequently DNA was stretched to a required force and the fluctuations of each bead were recorded simultaneously at *f*_*s*_ = 78.125kHz for a period of 6 seconds. These measurements were performed at DNA tensions varying between lpN and 5pN with 0.5 pN incremental steps, with 3 repeats per each force.

Prior to the measurements the laser power was adjusted to reach the nominal trap stiffness of *κ* _1_ = *κ* _2_ 0.05pN for each optical trap.

Data were acquired using a custom-made automation script. High-frequency force data and the distance between the beads measured at ≈50Hz were saved as an h5 file and further processed using Matlab.

The final numbers of molecules used for each length and each buffer condition are as in Table.II

For each single molecule experiment, we corrected forces according to Supplementary Information (section S6) to account for small positive forces – in the absence of DNA – between beads due to the cross-talk between traps at small trap separations.

### 4. Data processing

Raw data in the form of force signals from both traps were used to extract the DNA length fluctuations as follows: at each time point the length of the DNA molecule was calculated as

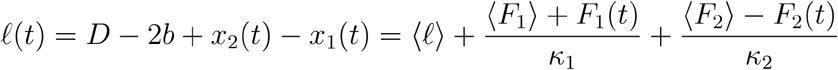

where *D* is the distance between trap centres, *b* the radius of each bead, ⟨F_2_⟩ = − ⟨F_1_⟩ = *F* Fare the mean force in each trap averaged over the measurement period (6 seconds), where *F* is the magnitude of the desired force, *κ*_1_, *κ*_2_ are the trap stiffness values for trap 1 and trap 2, and are nominally equal and ⟨ℓ⟩ is the mean DNA length, calculated as the distance between the beads averaged over the measurement period (6 seconds) with 2b subtracted, and *F*_1_, and *F*_2_ are the instantaneous forces on the beads in trap 1 and trap 2.

The power spectrum *P*_ℓ_*(ω)* of the time-series of DNA length ℓ *(t)* is then calculated by standard Fast Fourier Transform **(FFT)** in MatLab,

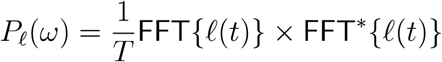

where * indicates complex conjugate and *T* = 5000msec is the maximum observation time in the data. The power spectrum is then block-averaged on a log-frequency scale [6] with *n* = 25 blocks, to reduce the noise on the raw power spectrum estimate. The zero-frequency (DC) component is removed and only frequencies less than the Nyquist frequency (*f*_*s*_/2) are retained. The velocity power spectrum is then calculated by *P*_*ν*_ (*ω*) = *ω*^2^ *P*_ℓ_ (*ω*).

As described in the main text, the velocity power spectrum is then fit with Eqn.S18 (Supplementary Information) at each force *F* to obtain *ζ* (*F*), *κ* (*F*) and *m*_*eff*_*(F)*.

## Notes

### Competing Interest Statement

The authors have declared no competing interest.

### Summary of Updates

Changes to abstract, introduction and discussion to emphasise unexpected nature of intramolecular friction within DNA (there are no obvious large energy barriers), the implications for molecular dynamics simulations of DNA and the inclusion of more detail on the derivation of main theoretical result.

### Funding

- Engineering and Physical Sciences Research Council: EP/R014604/1
- The Francis Crick Institute: FC001750

