## Supplementary figures and derivations for "Bending friction: a new mechanism of dissipation within DNA explains its slow looping dynamics"

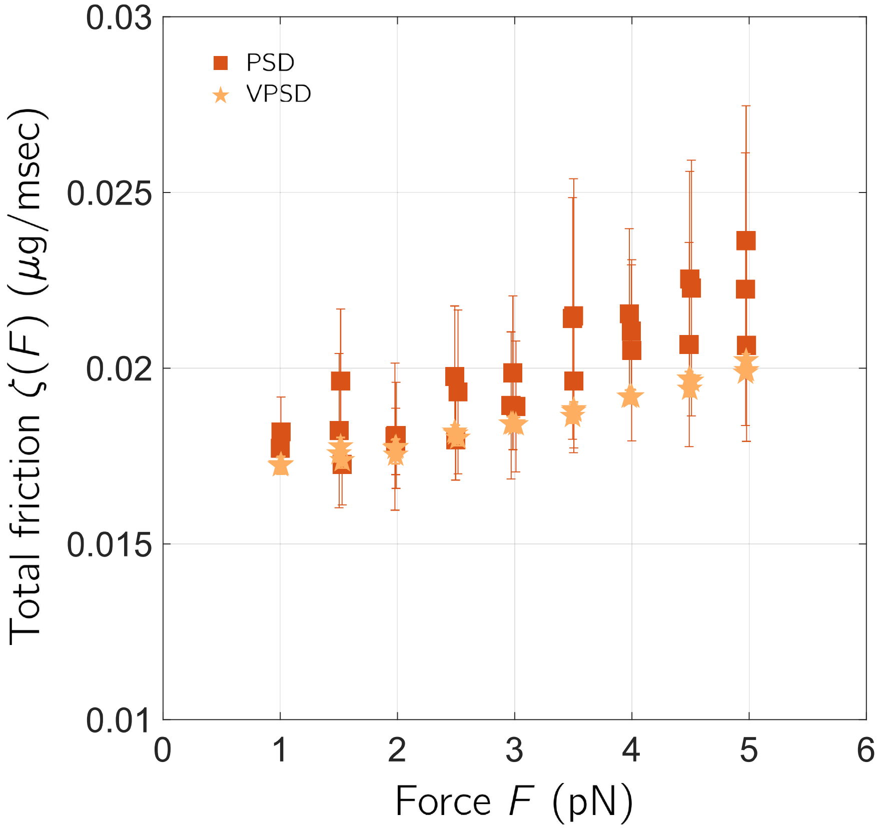


FIG. S1. Comparison of inferred friction of system + DNA using the velocity power spectrum (closed orange pentagrams) vs the power spectrum (closed red squares) for the same 2.7kbp DNA molecule. The model of the power spectrum does not include the high-frequency corrections since there is not enough information in the tail to determine the effective mass and the fitting to the PSD is performed between 0.1kHz and 15kHz to avoid increasing noise in power spectrum estimates at low frequency. We see using the power spectrum produces an estimate of friction which is more noisy and with larger error bars, but consistent with the more precise estimate using the VPSD. Error bars are 67% confidence intervals on parameter value estimates from non-linear regression.

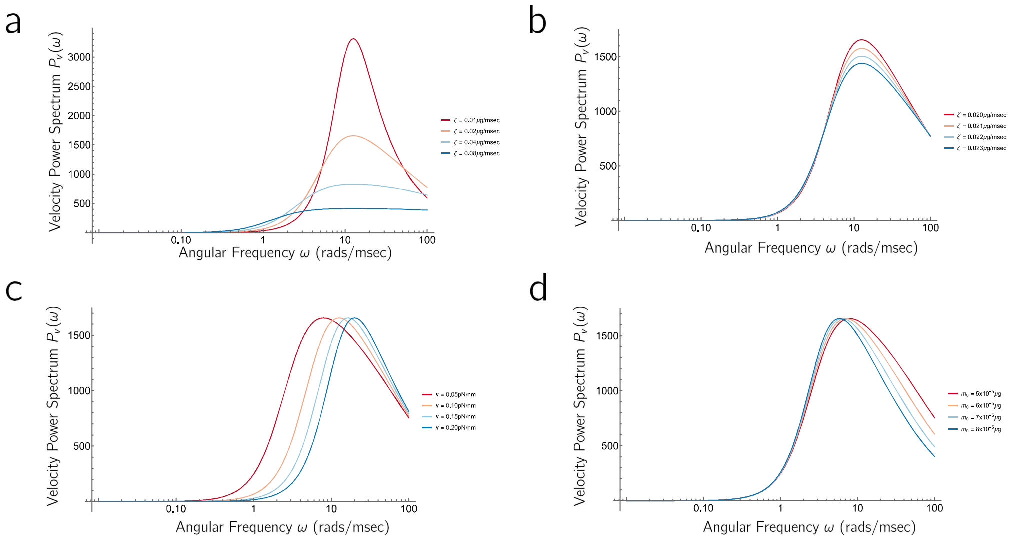


FIG. S2. **Maximum of VPSD is inversely proportional to friction constant.** To demonstrate the sensitivity of the VPSD to changing friction constant (of DNA), we can plot the VPSD in Eqn.S18 (Supplementary Information) for various values of total friction $\zeta$, elastic constant $\kappa$ and effective hydrodynamics mass $m_0$ for values corresponding to those determined by experiment on 2.6 kbp DNA. **a)** we plot the VPSD for different values of friction $\zeta = \{0.01, 0.02, 0.04, 0.08\}$ $\mu$g/msec, $\kappa = 0.1$ pN/nm and $m_0 = 5 \times 10^{-5}$ $\mu$g — which corresponds to approximately to the value of $m_0$ obtained by fitting the VPSD for 2.6 kbp DNA, as shown in Fig.S4. We see a dramatic drop in peak height for increasing friction, showing that the maximum of the VPSD is inversely proportional to the friction constant. **b)** Plots of the VPSD for a range of $\zeta$ more representative of the values determined from experiment by stretching DNA between $F = 1$ pN and $F = 5$ pN, $0.02 \leq \zeta \leq 0.023$ $\mu$g/msec, then we the same behaviour, but with a similar change in peak height observed empirically in the VPSD plotted in the main text in Fig.2b. **c)** To check that a decreasing peak height cannot be obtained by changing $\kappa$ (or $m_0$ in part d of this figure), we plot for a fixed value of $\zeta = 0.02$ $\mu$g/msec, the VPSD for $\kappa = \{0.05, 0.10, 0.15, 0.20\}$ pN/nm, which corresponds to the range of $\kappa$ found for stretching 2.6 kbp DNA between $F = 1$ pN and $F = 5$ pN; we see that the peak height does not change, but moves to higher frequencies for increasing $\kappa$. This explains why in the empirical VPSD (Fig.2b, main text), as the tension $F$ increases, the peak decreases in height and also moves to higher frequencies, as both the $\zeta$ and $\kappa$ increase with increasing tension, due to the non-linear frictional and elastic response of DNA. **d)** Plot of the VPSD for $m_0 = \{5 \times 10^{-5}, 6 \times 10^{-5}, 7 \times 10^{-5}, 8 \times 10^{-5}\}$ $\mu$g —which is a range which far exceeds the values found from fits in Fig.S4, which varies between $5.5 \times 10^{-5}$ $\mu$g and $6 \times 10^{-5}$ $\mu$g — for $\zeta = 0.02$ $\mu$g/msec and $\kappa = 0.05$ pN/nm and we see that again the peak position decreases, moderately, for increasing $m_0$ and is not the cause of the decreasing peak height.

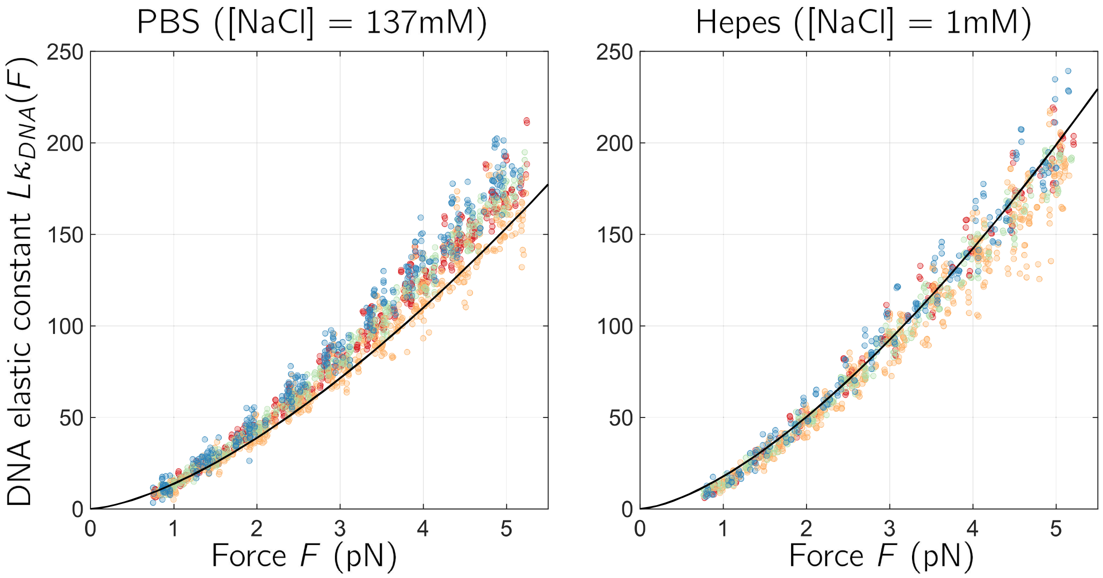


FIG. S3. Elastic constant derived from fluctuations is consistent with WLC fits of force-extension fits for normal PBS buffer and low salt Hepes buffer. Different colours correspond to normalised elastic constant derived from fitting the VPSD, with different contour lengths of DNA: 2.7 kbp ($L$ = 902.7 nm; red circles), 4.5kbp ($L$ = 1530.0 nm;yellow/orange circles), 6.5 kbp ($L$ = 2200.8nm;green circles), 8.8 kbp ($L$ = 2997.1 nm;blue circles). The solid line is a plot of the elastic constant of an extensible WLC $L\kappa_{\Delta R}(F) = \frac{4\sqrt{\kappa_B}F^{3/2}}{k_BT}$ (Supplementary Information: Eqn.S29) with persistence length $\ell_P$ = 47.9 nm for PBS and $\ell_P$ = 81.9 nm, which are derived from force-extension fits of an extensible WLC, but where here we do not need to consider backbone stretching terms at these small forces (see Supplementary Information: section S3). The number of single molecules used are as described in table.II in the Methods.

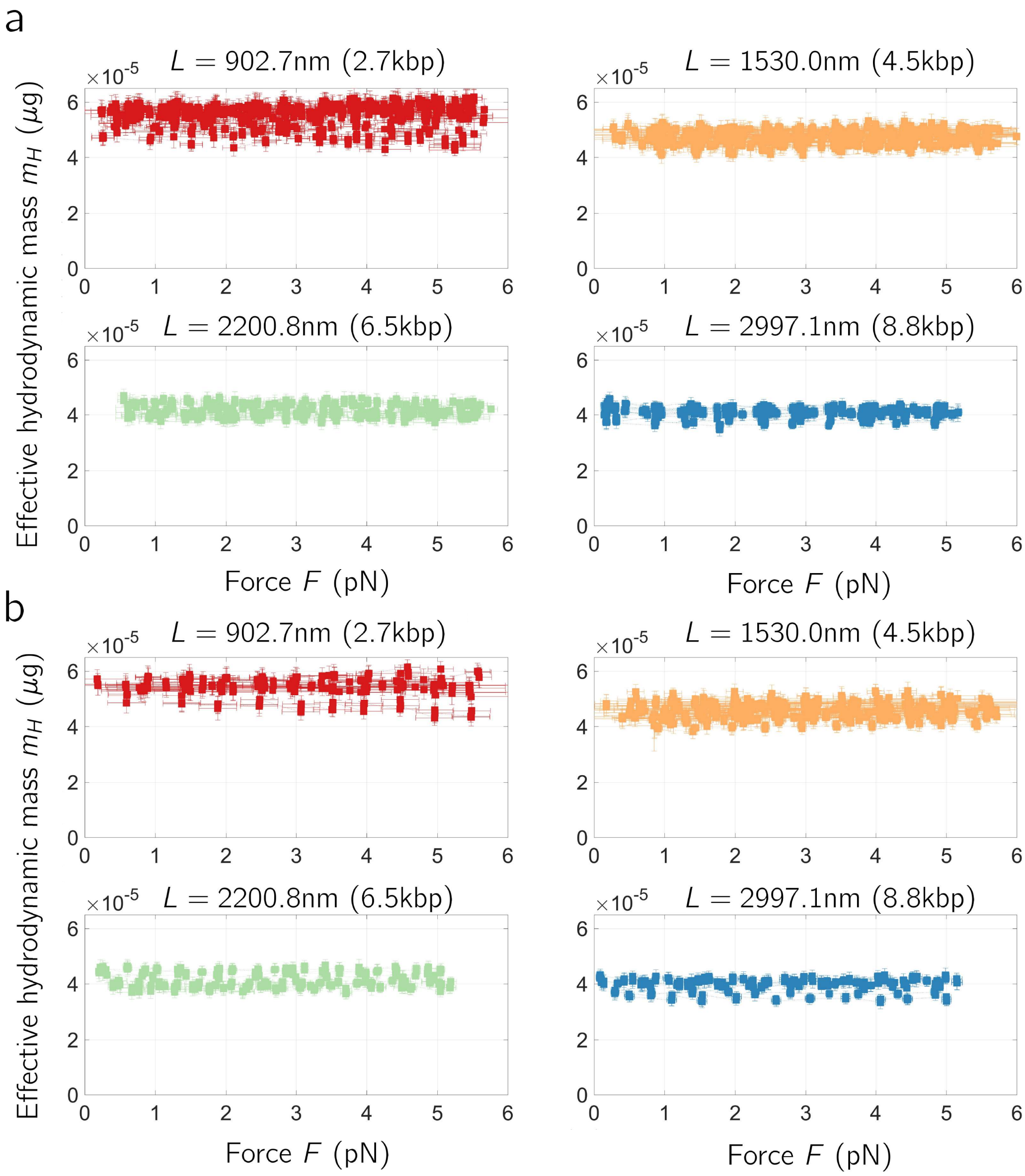


FIG. S4. Effective hydrodynamic mass from fitting to VPSD with DNA as function of force is independent of force. This indicates that there are no artefacts that are effecting estimation of the friction constant from VPSD, and is further rationalised, since in Extended Figure 2, we see the signal of changing friction and changing effective mass are very different. a) Normal salt PBS buffer and b) low-salt Hepes buffer. Error bars are 67% confidence intervals on parameter value estimates from non-linear regression.

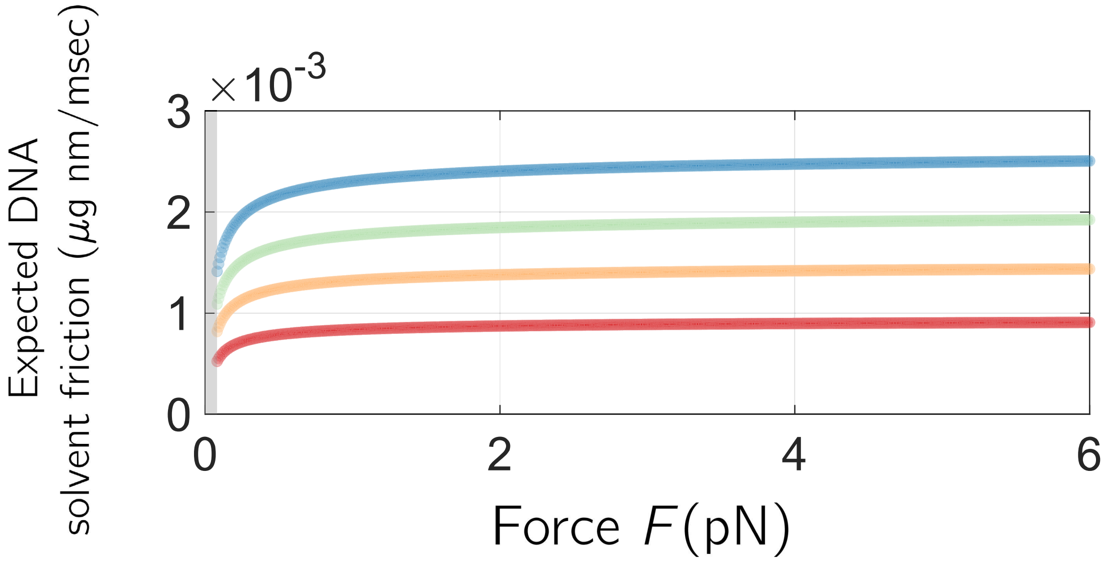


FIG. S5. The expected excess DNA friction, if it were due to solvent dissipation assuming DNA is highly stretched ($F > 0.08$pN) using $\zeta_s = 2\pi\eta\ell(F)/\ln(\ell(F)/w)$, where $\ell(F) = L\left(1 - \sqrt{\frac{k_BT}{4F\ell_P}}\right)$ is the extension of the WLC as a function of force $F$. This is in stark contrast to the measured excess DNA friction which increases linearly with force and increases with decreasing contour length (Main text Fig.3)

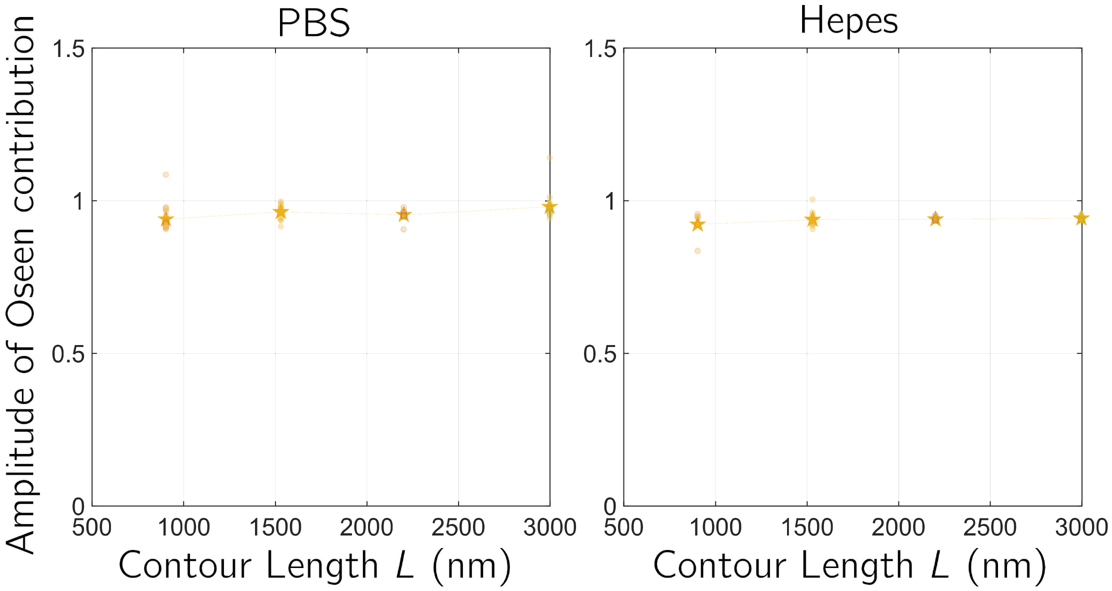


FIG. S6. Values of the fitting parameter $A$ used to subtract off background bead and solvent friction hydrodynamics. We determined the excess friction of DNA as the residual friction over and above the background bead friction and DNA solvent friction, from each single molecule measurement of total friction $\zeta(F)$, by fitting to the total friction as $\zeta(F) = \Delta\zeta_{DNA}(F) + A\zeta_{background}(r(F))$, using Eqn.5 in the main text, where the second term is the effective bead friction due to Oseen flows measured without DNA for the same beads used to do the measurements with DNA, and $r(F) = \ell(F) + 2b$, is the distance between bead centres, where $\ell(F) = L(1 - \sqrt{k_B T/4F\ell_p})$ is the force-dependent equilibrium end-to-end length of DNA. $A$ is a fitting parameter, for each molecule, and represents the additional effects due to solvent dissipation of DNA. Error bars are 67% confidence intervals on parameter value estimates from non-linear regression. The values of the fitting parameter are close to one, as one would expect, although in total the values tend to be less than one, which could indicate more complicated hydrodynamic interactions between the beads and DNA than a simple additive model would suggest. This effect is reminiscent of turbulent drag reduction[22], although this phenomenon would only play a role at relatively high Reynold's number, which is not the case here.

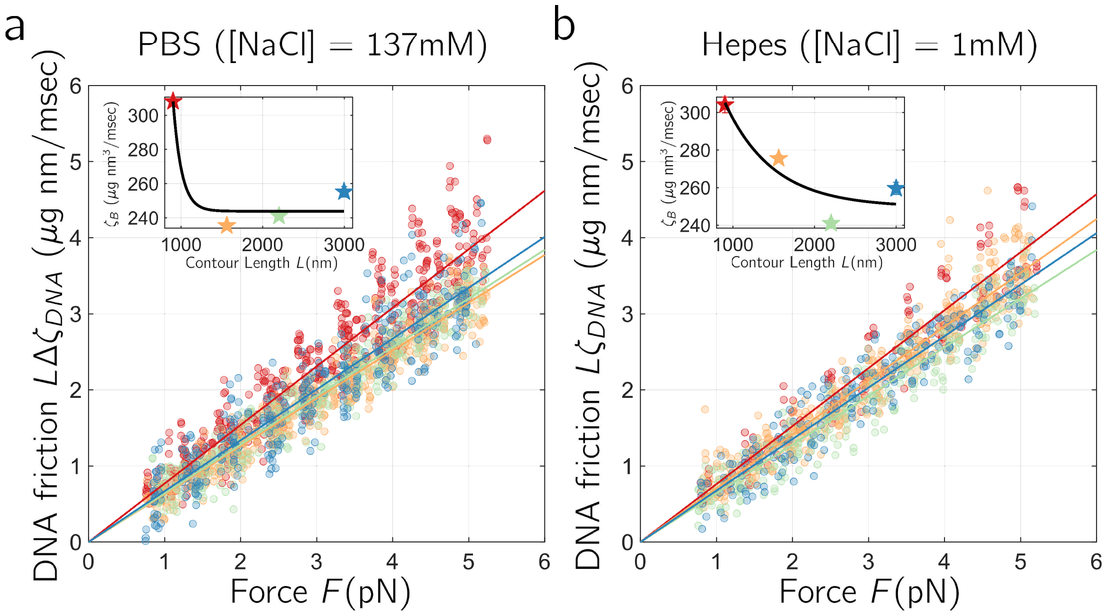


FIG. S7. Plot and fits of normalised excess DNA friction $L\Delta\zeta_{DNA}$ vs force $F$, showing good data collapse. a) is the data for normal salt PBS buffer, while b) is for the low-salt Hepes buffer, where different colours correspond to different contour lengths, with the length as indicated in the inset plots. Fits to each molecule are using Eqn.5 in the main text with single fit parameter $\zeta_B$, the mean and standard error over all molecules is plotted in the inset as a function of contour length $L$. We see the parameter $\zeta_B$ increases for decreasing contour length, which we attribute to finite sized effects, as has been previously used to explain the apparent length dependence for the persistence length of DNA[20]. For each salt condition, we fit with an exponentially saturating function and determine $\zeta_B$ as the infinite-length or plateau value. This gives $\zeta_B = 238 \pm 20$ $\mu$g nm$^3$/ms for the PBS buffer and $\zeta_B = 249 \pm 32$ $\mu$g nm$^3$/ms, for the Hepes buffer (67% confidence intervals on parameter value estimates from non-linear regression).

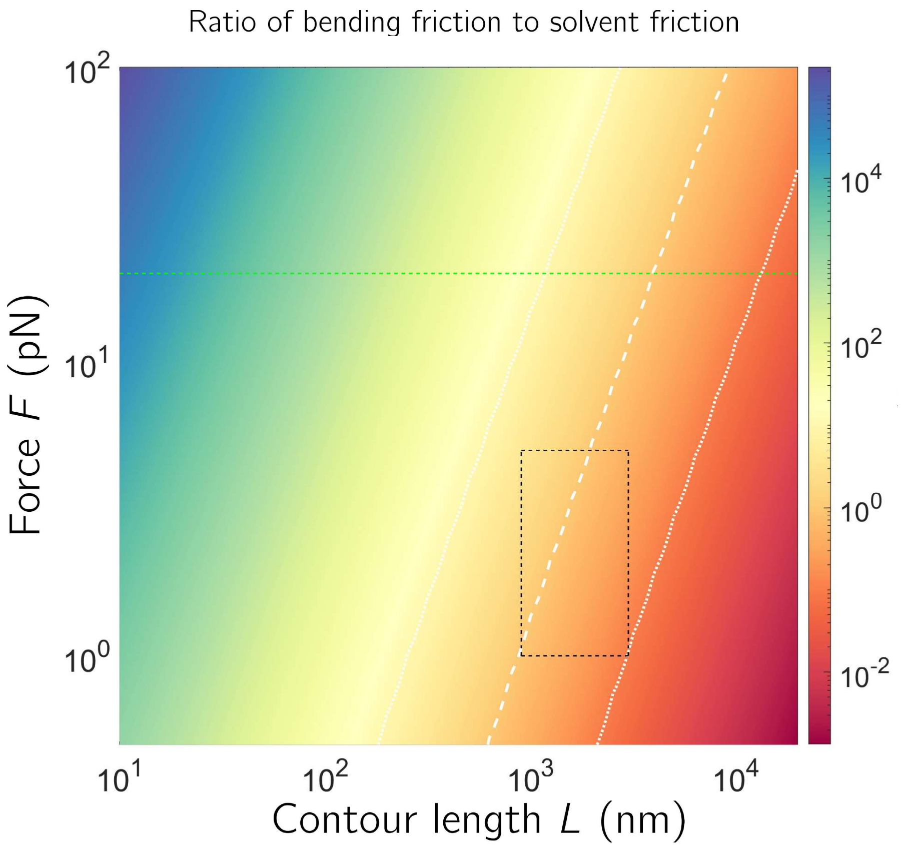


FIG. S8. A heat map of the ratio of bending friction to solvent friction, as as function of force $F$ and contour length $L$, given our measurement of $\zeta_B = 241$ $\mu$gnm$^3$/ms, using Eqn.5 in the main text and the expected solvent friction assuming DNA is completely stretched and width $w = 2.4$ nm ($\zeta_s \approx \frac{2\pi\eta L}{\ln(L/w)}$). The white dashed line indicates where this ratio =1, the dotted lines values of 10 and 0.1, the green dotted line indicates $F = 20$pN (which have been observed within cells in vivo [16, 19]) the black dashed box is the range of forces and contour lengths probed in the experiments in this paper. This plot exemplifies the interplay between tension and chain length on the sources of dissipation within DNA, where for DNA shorter than a few persistence lengths bending dissipation will always dominate, and also for longer lengths, at sufficiently large forces. Note that this calculation and figure is only valid at high stretch, in excess of the critical force $k_BT/\ell_P \approx 0.08$ pN, and that a direct to comparison to Fig.4a of the relative contribution of bending vs solvent friction for each mode at zero force, is not possible, since the latter ignores the contributions to solvent friction of long range hydrodynamics.

### S1. THE VELOCITY POWER SPECTRUM OF BEAD SYSTEM FLUCTUATIONS

Using the property of Fourier Transforms, the relationship between the power spectrum of a coordinate $x(t)$ and of the velocity $\dot{x}(t)$ is

$$P_{\dot{x}}(\omega) = \omega^2 P_x(\omega). \tag{S1}$$

Below we will calculate the theoretical power spectrum and for each case the velocity power spectrum can be calculated using this scaling.

#### Power spectrum of a single bead

We start with the standard description of the power spectrum of a single bead of radius $b$ in the overdamped, but non-stationary regime [11]:

$$P(\omega) = \frac{4k_B T\tilde{\zeta}(\omega)}{\left(\kappa - m_H(\omega)\omega^2\right)^2 + \tilde{\zeta}^2(\omega)\omega^2}, \tag{S2}$$

with a frequency dependent effective friction $\tilde{\zeta}(\omega) = \zeta_0\left(1 + b/\xi(\omega)\right)$ and frequency dependent hydrodynamic mass is $m_H(\omega) = 3\pi b^2\rho\xi(\omega) + \frac{2}{3}\pi b^3\rho$, where $\zeta_0 = 6\pi\eta b$ is the normal (zero frequency) Stokes friction of the bead, and fluctuations at a given frequency $\omega$ have an effective penetration depth $\xi = \sqrt{\frac{2\eta}{\rho\omega}}$, where $\rho \approx 10^{-15}\mu\text{g/nm}^3$ is the density of water and $\eta \approx 10^{-6}\mu\text{g/msec/nm}$. Physically, the penetration depth corresponds to the depth of fluid at the beads surface that is effectively entrained as it cannot respond quickly enough for fluctuations at a given frequency. So the additional friction friction at finite frequency arises due to friction between this entrained fluid and the stationary fluid beyond. The hydrodynamic mass term is the effective mass of the entrained fluid and of the displaced fluid, respectively.

**Power spectrum of two beads with stationary flows**

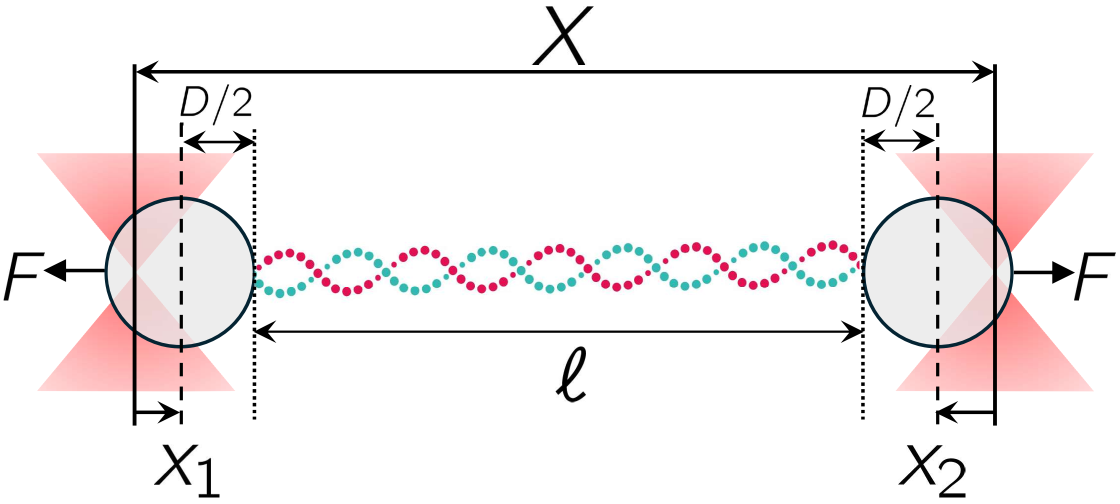


FIG. S9.

In Fig.S9, we have a schematic of the dual-trap and bead optical trap system. We assume the trap centres are placed a distance $D$ apart, and the displacement of each bead centre from the centre of each trap is denoted by $x_1$ and $x_2$, respectively (in the figure $x_1 > 0$ and $x_2 < 0$, assuming positive distances are left to right).

With two beads, we need to consider the interactions between beads, which we assume is mediated by the Oseen tensor $\mathsf{L}$ which is a symmetric matrix with elements $L_{ij}$, where the diagonal elements represent interactions of each bead with the solvent and the off-diagonal are the interactions between beads, mediated by the fluid hydrodynamics, at a separation $r$ between centres of the beads. These matrix elements are given by:

$$L_{11} = L_{22} = \frac{1}{6\pi\eta b} \tag{S3}$$

$$L_{12} = L_{21} = \frac{1}{4\pi\eta r}. \tag{S4}$$

The Fourier transform of the Langevin equation for bead positions $x_1(t)$ and $x_2(t)$ is then in vectorial form:

$$i\omega \boldsymbol{X} = \mathsf{L}\left(-\kappa \mathsf{I} \boldsymbol{X}(\omega) + \boldsymbol{F}(\omega)\right), \tag{S5}$$

where

$$\boldsymbol{X}(\omega) = \begin{pmatrix} X_1(\omega) \\ X_2(\omega) \end{pmatrix},$$

&

$$\boldsymbol{F}(\omega) = \begin{pmatrix} F_1(\omega) \\ F_2(\omega) \end{pmatrix},$$

with $X_j(\omega) = \int_{-\infty}^{\infty} \mathrm{d}t x_j(t) e^{-i\omega t}$ and $F_j(\omega) = \int_{-\infty}^{\infty} \mathrm{d}t f_j(t) e^{-i\omega t}$, where $f_1(t)$ and $f_2(t)$ are the Langevin noise terms acting on each bead, $\kappa$ is the spring constant of each trap and $\mathsf{I}$ is the identity matrix. This equation can be simply re-arranged and inverted to give the solution

$$\boldsymbol{X(\omega)} = \mathsf{J}(\omega)\boldsymbol{F}(\omega),$$

where $\mathsf{J}(\omega)$ is a compliance matrix given by

$$\mathsf{J}(\omega) = \left(i\omega \mathsf{L}^{-1} + \kappa \mathsf{I}\right)^{-1}$$

and the inverse of the Oseen tensor is

$$\mathsf{L}^{-1} = \begin{pmatrix} \zeta_{11} & -\zeta_{12} \\ -\zeta_{21} & \zeta_{22} \end{pmatrix},$$

where

$$\zeta_{11} = \zeta_{22} = \frac{L_{11}}{L_{11}^2 - L_{12}^2} = \frac{6\pi\eta b}{1-\epsilon^2},$$

and

$$\zeta_{12} = \zeta_{21} = \frac{L_{12}}{L_{11}^2 - L_{12}^2} = \frac{4\pi\eta r\epsilon^2}{1-\epsilon^2},$$

where $\epsilon = \frac{3b}{2r}$; in the limit that $b/r \sim \epsilon \to 0$, $\zeta_{11} \to 6\pi\eta b$ and $\zeta_{12} \to 0$, as we would expect. Using this and $\mathsf{J}(\omega)$, we can calculate the power spectrum of the difference in bead positions $\Delta x(t) = x_2(t) - x_1(t)$:

$$\begin{aligned}
P_{\Delta x}(\omega) &= \langle |X_2(\omega) - X_1(\omega)|^2 \rangle && \text{(S6)}\\
&= \langle (X_2(\omega) - X_1(\omega))(X_2(\omega) - X_1(\omega))^* \rangle && \text{(S7)}\\
&= \langle |X_1(\omega)|^2 \rangle + \langle |X_2(\omega)|^2 \rangle - 2\mathsf{Re}(\langle X_1(\omega) X_2^*(\omega) \rangle) && \text{(S8)}\\
&= P_1(\omega) + P_2(\omega) - 2P_{12}(\omega) && \text{(S9)}\\
&= -\frac{8k_BT}{\omega}\left(\mathsf{Im}(J_{11}) - \mathsf{Im}(J_{12})\right), && \text{(S10)}
\end{aligned}$$

where have used the fact that the cross-power spectra between beads $P_{12}(\omega) = \mathsf{Re}(\langle X_1 X_2^* \rangle)$ — which can be shown, straightforwardly, from its definition $P_{12}(\omega) = \int_{-\infty}^{\infty} \mathrm{d}t \langle x_1(0) x_2(t) \rangle e^{-i\omega t}$ — and the Fluctuation-Dissipation Theorem (FDT), where in general, $P(\omega) = -\frac{4k_BT}{\omega}\mathsf{Im}(J)$, where we assume the PSD is single-sided (i.e. that power from negative frequencies are folded over to positive frequencies to give double the PSD compared to the double-sided PSD). After some computation the power spectrum is given by

$$P_{\Delta x}(\omega) = \frac{8k_BT(\zeta_{11} + \zeta_{12})}{\kappa^2 + (\zeta_{11} + \zeta_{12})\omega^2},$$

where we can obtain a simple expression for the total friction:

$$\zeta_{11} + \zeta_{12} = \frac{12\pi\eta b r}{2r - 3b}, \tag{S11}$$

which is the expression we use in the main text to fit the $r$ dependence of the background friction without DNA attached.

#### Power spectrum of two beads with non-stationary flows

In this section we repeat the calculation for calculation of $P_{\Delta x}(\omega)$, but include the non-stationary flows around each bead, but ignore the analogous high frequency corrections to hydrodynamic interaction between beads[1].

Including the non-stationary/frequency dependent terms for the fluctuations around each bead, we obtain

$$L_{11}(\omega) = L_{22}(\omega) = \frac{1}{\zeta_0(1 + \frac{b}{\xi(\omega)}) + i\omega m_H(\omega)} \tag{S12}$$

where $\zeta_0 = 6\pi\eta b$ and $L_{12}$ is unchanged. The calculation carries through the same as above, but with the frequency dependent form of the Oseen tensor given, the expression for $P_{\Delta x}(\omega)$ does not give a simple expression that is amenable to extracting the friction of DNA. We now show that the frequency dependent dynamics can be simplified for the frequency regime relevant to our experiments, where the maximum frequency we analyse the data is at 15kHz, which when combined with the friction and elastic constant of DNA, gives a single total friction and elastic constant, as well as an effective mass term related to the entrained fluid around each bead.

In general, $\zeta_{11}(\omega)$ and $\zeta_{12}(\omega)$ will be complex, where the real parts will contribute to dissipation and the imaginary parts giving rise to terms related to the effective mass of fluid entrained around each bead. Hence, $\mathsf{Re}(\zeta_{11}(\omega) + \zeta_{12}(\omega))$ will play an analogous role to the total friction, calculated above in Eqn.S11, but with frequency dependent terms. The expression for $\mathsf{Re}(\zeta_{11}(\omega) + \zeta_{12}(\omega))$ is complicated, but we evaluate it and plot it vs frequency in Fig.S10, for $r = 2770$nm and $r = 4557$nm, corresponding to the expected distance between bead centres for 2.6kb ($L \approx 900$nm) and 8.8kb ($L \approx 3000$nm), stretched at $F = 1$pN, which is the minimum force we use in our experiments. We see that up to up to 15kHz ($\approx$ 95radians/msec) the dependence on $\omega$ is relatively weak, and so we can approximately treat it as a constant, with the expectation that we may end up overestimating the zero frequency friction. We see the asymptote for $\omega \to 0$ agrees with the dashed lines, which is Eqn.S11, the expected frequency independent friction.

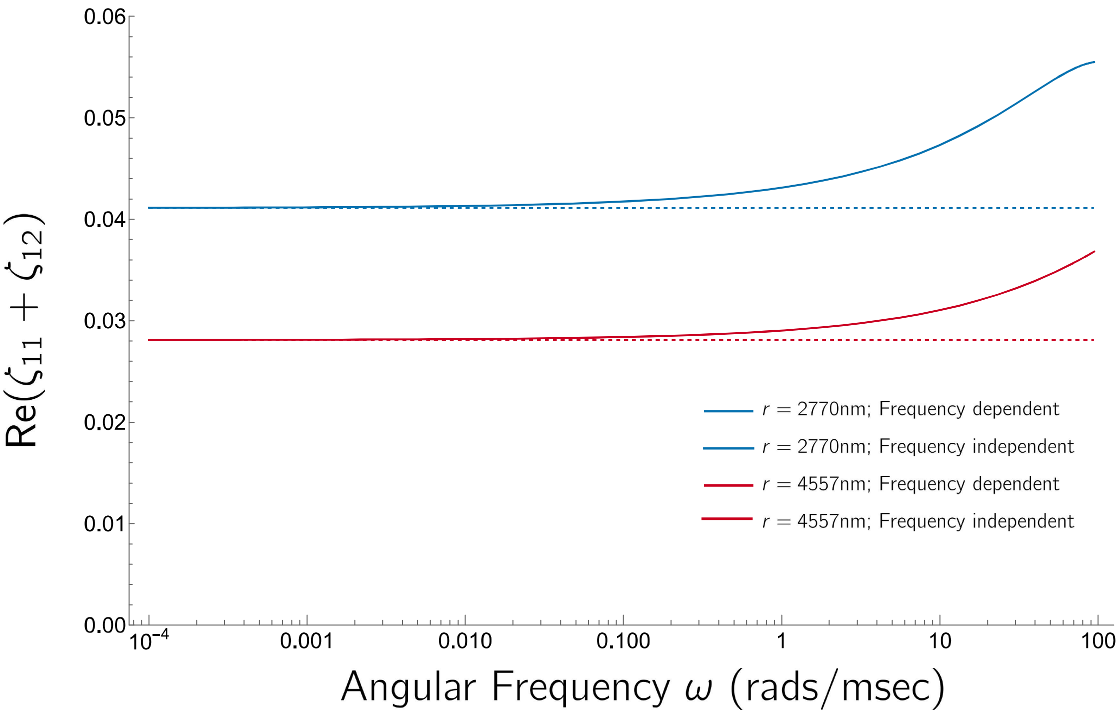


FIG. S10.

On the other hand, in Fig.S11 we plot the effective mass $m_H(\omega) = \frac{1}{\omega}\mathsf{Im}(\zeta_{11}(\omega) + \zeta_{12}(\omega))$ — on a log-log scale — and we see has a much stronger frequency dependence which cannot be ignored, and is very well approximated by $\frac{1}{\sqrt{\omega}}$. Again, the exact expression for $\frac{1}{\omega}\mathsf{Im}(\zeta_{11}(\omega) + \zeta_{12}(\omega))$ is complicated, but the leading order term of the series expansion is

$$m_H(\omega) = \frac{1}{\omega}\mathsf{Im}(\zeta_{11}(\omega) + \zeta_{12}(\omega)) \approx \frac{12\pi r^2 b^3 \rho \sqrt{\frac{2\eta}{\rho\omega}}}{(2r - 3b)^2} = m_0 \frac{\xi(\omega)}{b} \tag{S13}$$

where $m_0 = \frac{12\pi r^2 b^3 \rho}{(2r-3b)^2}$. Although, there is an $r$-dependence to this effective mass term, it is weak over the range of separations used in the experiments, as the chain is at high stretch close to its contour length, or in limit when $r \gg b$, where $m_0$ is independent of $r$.

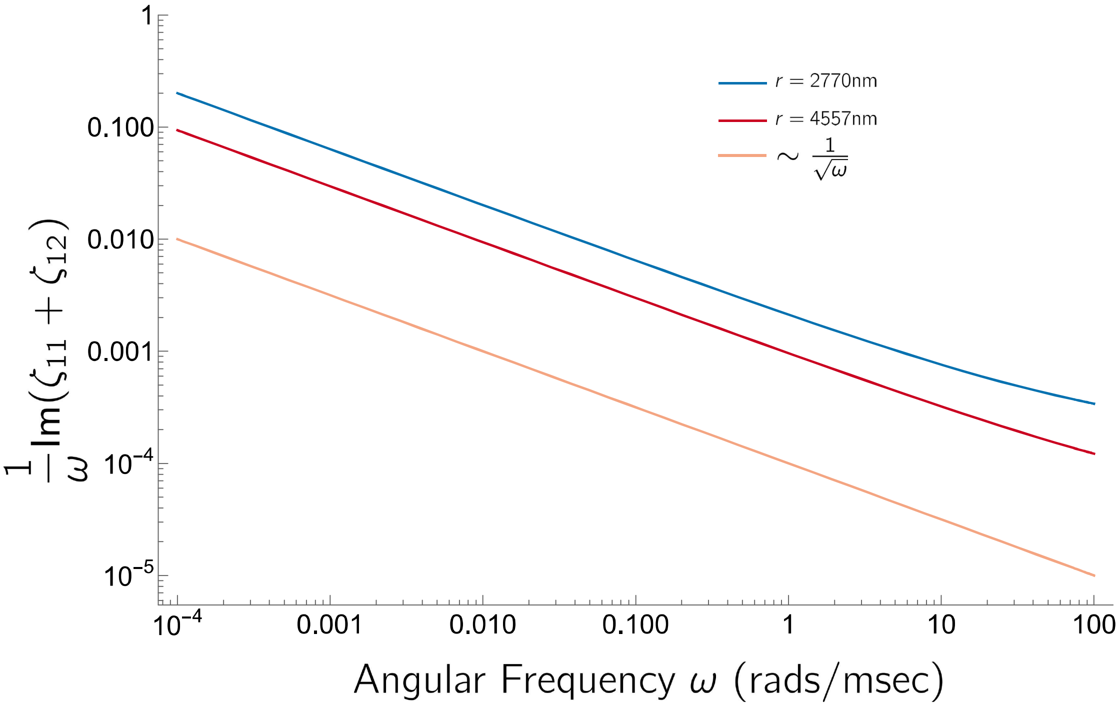


FIG. S11.

Making these approximation that $\mathsf{Re}(\zeta_{11}(\omega) + \zeta_{12}(\omega))$ is roughly independent of frequency, and that $\frac{1}{\omega}\mathsf{Im}(\zeta_{11}(\omega) + \zeta_{12}(\omega)) \sim \frac{m_0}{\sqrt{\omega}}$, after some calculation we arrive at an approximate expression for the power spectrum of the bead separations.

$$P_{\Delta x}(\omega) = \frac{8k_BT(\zeta_{11}+\zeta_{22})}{(\kappa - m_H(\omega)\omega^2)^2 + (\zeta_{11}+\zeta_{22})^2\omega^2}, \tag{S14}$$

where implicitly the $\zeta$'s without the $\omega$-dependence shown indicate use of the zero-frequency expression in Eqn.S11.

**Power Spectrum of two beads with DNA with non-stationary flow approximations**

Having established the relevant frictional forces for a two beads due to hydrodynamic interactions for the frequencies considered in the experiment, we incorporate the friction (& elasticity) of DNA when tethered between the two beads.

The elastic forces follow from writing a potential $U(x_1, x_2)$:

$$U(x_1,x_2) = \frac{1}{2}\kappa x_1^2 + \frac{1}{2}\kappa x_1^2 + \frac{1}{2}\kappa_{DNA}\ell^2(x_1,x_2),$$

where in reference to Fig.S9 $\ell = D + (x_2 - x_1) - 2b$, $\kappa$ is the trap spring constant and $\kappa_{DNA}$ for (non-linear) elasticity of the DNA molecule. The elastic forces on each bead are then

$$-\frac{\partial U}{\partial x_1} = -\kappa x_1 + \kappa_{DNA}\ell = -\kappa x_1 + \kappa_{DNA}(x_2 - x_1) + F_0$$
$$-\frac{\partial U}{\partial x_2} = -\kappa x_2 - \kappa_{DNA}\ell = -\kappa x_2 - \kappa_{DNA}(x_2 - x_1) - F_0,$$

where $F_0 = \kappa_{DNA}(D - 2b)$ is the fixed force that ensures the the DNA molecule has separation $\ell$. From here on we ignore this term as it has no effect on the dynamics of $x_1$ and $x_2$. Analogously, we can calculate the frictional forces, by using the Rayleigh dissipation function $\mathcal{D}(\dot{x}_1, \dot{x}_2)$:

$$\mathcal{D}(\dot{x}_1,\dot{x}_2) = \frac{1}{2}\zeta_{11}\dot{x}_1^2 + \frac{1}{2}\zeta_{22}\dot{x}_1^2 - \frac{1}{2}\zeta_{12}\dot{x}_1\dot{x}_2 - \frac{1}{2}\zeta_{21}\dot{x}_2\dot{x}_1 + \frac{1}{2}\zeta_{DNA}\dot{\ell}^2.$$

Using, the symmetry of the friction coefficients $\zeta_{11} = \zeta_{22}$ and $\zeta_{12} = \zeta_{21}$, the frictional forces are

$$
\begin{aligned}
-\frac{\partial \mathcal{D}}{\partial \dot{x}_1} &= -\zeta_{11}\dot{x}_1 + \zeta_{12}\dot{x}_2 + \zeta_{DNA}(\dot{x}_2 - \dot{x}_1) \\
-\frac{\partial \mathcal{D}}{\partial \dot{x}_2} &= -\zeta_{11}\dot{x}_2 + \zeta_{12}\dot{x}_1 - \zeta_{DNA}(\dot{x}_2 - \dot{x}_1).
\end{aligned}
$$

The force balance on both beads then becomes the following matrix equation:

$$\mathsf{L}^{-1}\dot{\boldsymbol{x}} + \mathsf{K}\boldsymbol{x} = \boldsymbol{f},$$

where $\boldsymbol{x} = (x_1, x_2)^T$, the inverse mobility matrix

$$\mathsf{L}^{-1} = \begin{pmatrix} \zeta_{11} + \zeta_{DNA} & -(\zeta_{12} + \zeta_{DNA}) \\ -(\zeta_{12} + \zeta_{DNA}) & \zeta_{11} + \zeta_{DNA} \end{pmatrix},$$

$$\mathsf{K} = \begin{pmatrix} \kappa + \kappa_{DNA} & -\kappa_{DNA} \\ -\kappa_{DNA} & \kappa + \kappa_{DNA} \end{pmatrix},$$

and where $\boldsymbol{f} = (f_1, f_2)^T$ are the Langevin noise force terms with zero mean and correlation:

$$\langle f_i(t) f_j(t') \rangle = 2k_B T (\mathsf{L}^{-1})_{ij} \delta(t - t').$$

From this modified inverse mobility matrix, we can then follow through the procedures of the previous sections to include non-stationary frequency dependent mass terms to calculate the PSD of the difference $\Delta x$ in trap positions as:

$$P_{\Delta x}(\omega) = \frac{4k_B T \zeta}{(\kappa - m_H(\omega)\omega^2)^2 + \zeta^2\omega^2}, \tag{S15}$$

where the total elastic constant is

$$\kappa = \kappa_{DNA} + \frac{1}{2}\kappa, \tag{S16}$$

and total friction constant is

$$\zeta = \zeta_{DNA} + \frac{1}{2}(\zeta_{11} + \zeta_{12}), \tag{S17}$$

where we have scaled the friction and elastic constants by $\frac{1}{2}$, which results in $4k_BT$ in the numerator instead of $8k_BT$ above. This results in an effective hydrodynamic mass of the the fluid entrained around each bead of

$$m_H(\omega) = \frac{m_0\xi(\omega)}{2b}.$$

Finally, using Eqn.S1, we obtain the velocity power spectrum (VPSD) used to fit the empirical VPSD from single molecule fluctuations in the main text:

$$P_{\Delta\dot{x}}(\omega) = \frac{4k_BT\zeta\omega^2}{\left(\kappa - m_H(\omega)\omega^2\right)^2 + \zeta^2\omega^2}. \tag{S18}$$

Note that for high frequency the VPSD decays with increasing $\omega$:

$$\lim_{\omega\to\infty}\{P_{\Delta\dot{x}}(\omega)\} \to \frac{4k_BT\zeta\omega^2}{m_H(\omega)\omega^4} = \frac{8k_BTb\zeta}{m_0\omega^{3/2}}\sqrt{\frac{\rho}{2\eta}},$$

which is as is observed in the experimentally determined VPSD, giving rise to a maximum.

### S2. SEMIFLEXIBLE POLYMER DYNAMICS WITH BENDING FRICTION: CALCULATING THE END-TO-END FRICTION UNDER TENSION

Semiflexible polymer dynamics are described by a stochastic partial differential equation describing the elastic, frictional and Brownian forces on each element of continuum rod, whose vectorial space curve is $\boldsymbol{R}(s,t)$, which is a function of the backbone coordinate $s$ and time $t$. Under tension, we assume that at high stretch ($F\ell_P \gg k_BT$) — or equivalently if $L \ll \ell_P$ — the space curve $\boldsymbol{R}(s,t) \approx \boldsymbol{R}_\perp$, the decomposition of the vector perpendicular to the direction of force. From here on and in the main text, we implicitly assume that $\boldsymbol{R}(s,t)$ corresponds to the perpendicular component. In this regime the elastic energy of the chain is

$$\mathcal{H}_e = \frac{1}{2}\kappa_B \int_0^L \mathrm{d}s \left(\frac{\partial^2 \boldsymbol{R}(s,t)}{\partial s^2}\right)^2 .$$

where $\kappa_B = k_BT\ell_p$ is the bending elastic constant. However, the external tension on the ends of the chain $\boldsymbol{F}$ means segments of the chain aligned with the direction of the applied force reduce the energy:

$$\mathcal{H}_F = -\int_0^L \mathrm{d}s \boldsymbol{u}\cdot\boldsymbol{F},$$

where $\boldsymbol{u} = \frac{\partial \boldsymbol{R}}{\partial s} \approx \boldsymbol{u}_\perp$ at high stretch, and so

$$\mathcal{H}_F \approx -F\int_0^L \mathrm{d}s\left(1 - \frac{1}{2}|\boldsymbol{u}_\perp|^2\right) = -FL + \frac{F}{2}\int_0^L \mathrm{d}s\left(\frac{\partial \boldsymbol{R}(s,t)}{\partial s}\right)^2 .$$

Hence the total energy is

$$\mathcal{H} = \mathcal{H}_e + \mathcal{H}_F = \frac{1}{2}\kappa_B \int_0^L \mathrm{d}s \left(\frac{\partial^2 \boldsymbol{R}(s,t)}{\partial s^2}\right)^2 + + \frac{F}{2}\int_0^L \mathrm{d}s\left(\frac{\partial \boldsymbol{R}(s,t)}{\partial s}\right)^2 , \quad \text{(S1)}$$

where we have ignored the constant energy term $FL$, as it does not depend on the shape of the chain. Local force balance on the chain then gives

$$-\tilde{\zeta}_s \frac{\partial \boldsymbol{R}(s,t)}{\partial t} - \frac{\delta \mathcal{H}}{\delta \boldsymbol{R}} + \boldsymbol{f}(s,t) = 0,$$

where the first term on the LHS is the local frictional force on the chain — with $\tilde{\zeta}_s$ is a solvent friction per unit length — the second term corresponds to the functional derivative of the Hamiltonian wrt to varying the space curve, which when added the local random Langevin force — the third term — according to force balance should equal zero on the RHS. As is standard in the field, the functional derivatives can be evaluated to give the equation for semiflexible polymer dynamics:

$$\tilde{\zeta}_s \frac{\partial \boldsymbol{R}(s,t)}{\partial t} - F \frac{\partial^2 \boldsymbol{R}(s,t)}{\partial s^2} + \kappa_B \frac{\partial^4 \boldsymbol{R}(s,t)}{\partial s^4} = \boldsymbol{f}(s,t) \tag{S2}$$

To include forces due to bending friction, we start with how much energy is dissipated due to local bending motion within a uniform rod with a bending friction constant $\zeta_B$. This is given by Rayleigh's dissipation function:

$$\mathcal{D} = \frac{1}{2}\zeta_B \int_0^L \mathrm{d}s \left( \frac{\partial}{\partial t} \frac{\partial^2 \boldsymbol{R}(s,t)}{\partial s^2} \right)^2 = \frac{1}{2}\zeta_B \int_0^L \mathrm{d}s \left( \frac{\partial^2 \dot{\boldsymbol{R}}(s,t)}{\partial s^2} \right)^2 .$$

This describes how dissipation due to bending friction is due the rate of change of local curvature, where the proportionality constant is the bending friction $\zeta_B$. The local frictional force is then given by the functional derivative of the dissipation function wrt to the local velocity of the space curve $\dot{\boldsymbol{R}}(s,t)$:

$$-\frac{\delta \mathcal{D}}{\delta \dot{\boldsymbol{R}}} = -\zeta_B \frac{\partial^4 \dot{\boldsymbol{R}}(s,t)}{\partial s^4}$$

Adding this to Eqn.S2 results in the modified Langevin equation which includes frictional forces due to bending dissipation (Eqn.1 in Methods):

$$\tilde{\zeta}_s \frac{\partial \boldsymbol{R}(s,t)}{\partial t} + \zeta_B \frac{\partial}{\partial t}\frac{\partial^4 \boldsymbol{R}(s,t)}{\partial s^4} - F\frac{\partial^2 \boldsymbol{R}(s,t)}{\partial s^2} + \kappa_B \frac{\partial^4 \boldsymbol{R}(s,t)}{\partial s^4} = \boldsymbol{f}(s,t).$$

Similar equations have appeared in the literature in different contexts [7, 17]. Now $\boldsymbol{f}(s,t)$ is a temporally white noise term, whose moments follow from the fluctuation dissipation theorem, but spatially coloured due the fact that dissipation occurs due to relative motion of adjacent points on the chain [10], which will show below in the normal mode analysis of Eqn.1.

**Normal mode analysis**

We can now analyse the dynamics of Eqn.1, by normal modes. Strictly, as Eqn.1 is a 4th order partial differential equation, the eigenfunction basis consists of combinations of sin, cos, sinh, cosh [23]. As DNA is attached with PEG linkers to each bead, the most appropriate boundary conditions at $s = 0$ and $s = L$ are there are no bending moments ($\partial_s^2 \boldsymbol{R}|_{s=0,s=L} = 0$) and no shear moments, which results in the following approximate relation for the wavenumber $q_n \approx \frac{1}{L}(n\pi + \frac{\pi}{2})$ for the $n^{th}$ mode, for $n > 0$. At high stretch, the fluctuations of the higher wavenumber modes will dominate and so we ignore the constant to give $q_n \approx \frac{\pi n}{L}$, and use the Fourier cosine eigenfunctions for simplicity.

The usual Fourier decomposition of the space curve then follows:

$$\boldsymbol{R}(s,t) = \sum_{n=0}^{\infty} \boldsymbol{r}_n(t) \cos(q_n s) \tag{S3}$$

$$\boldsymbol{r}_n(t) = \frac{1}{L}\int_0^L \mathrm{d}s \boldsymbol{R}(s,t) \cos(q_n s) \tag{S4}$$

Transforming Eqn.1 to a set of differential equations for the $\boldsymbol{r}_n(t)$ gives:

$$\zeta_n \frac{\mathrm{d}\boldsymbol{r}_n(t)}{\mathrm{d}t} + \kappa_n \boldsymbol{r}_n(t) = \boldsymbol{f}_n(t), \tag{S5}$$

where

$$\zeta_n = (\tilde{\zeta}_s + \zeta_B q_n^4)L \tag{S6}$$

$$\kappa_n = (\kappa_B q_n^4 + F q_n^2)L, \tag{S7}$$

where we have chosen to keep the contour length dependence in both these terms, so that the mode friction has a solvent friction proportional to $L$. The Langevin noise term for the $n^{th}$ mode is then:

$$\boldsymbol{f}_n(t) = \int_0^L \mathrm{d}s \boldsymbol{f}(s,t) \cos(q_n s),$$

with $2 \times 2$ correlation matrix:

$$\langle \boldsymbol{f}_n \boldsymbol{f}_m^T \rangle = 2 k_B T \zeta_n \delta_{nm} \mathbf{I},$$

where $\mathbf{I}$ is the $2 \times 2$ identity matrix, and $^T$ is the transpose of the column vector. This must be the form of the mode Langevin noise correlations, in order to ensure that the chain has the correct elastic properties in equilibrium. However, we can now use these to calculate the moments of the real-space Langevin noise

$$\boldsymbol{f}(s,t) = \frac{1}{L} \sum_{n=0}^{\infty} \boldsymbol{f}_n(t) \cos(q_n s).$$

As usual the mean is zero, but the 2nd moment can be evaluated to give

$$\begin{aligned}
\langle \boldsymbol{f}(s,t)\boldsymbol{f}^T(s',t')\rangle &= \frac{1}{L^2}\sum_{n=0}^{\infty}\sum_{m=0}^{\infty}\langle \boldsymbol{f}_n(t)\boldsymbol{f}_m^T(t')\rangle && \text{(S8)}\\
&= \frac{2k_BT}{L}\delta(t-t')\mathbf{I}\Bigg(\tilde{\zeta}_s\sum_{n=0}^{\infty}\cos(q_n s)\cos(q_n s') \\
&= \quad +\zeta_B\sum_{n=0}^{\infty}q_n^4\cos(q_n s)\cos(q_n s')\Bigg) && \text{(S9)}\\
&= 2k_BT\tilde{\zeta}_s\delta(t-t')\mathbf{I}\left(\tilde{\zeta}_s\delta(s-s')+\zeta_B\gamma_B(s-s')\right), && \text{(S10)}
\end{aligned}$$

where the first term represent the spatially white Langevin noise force due to solvent friction, while the second term represents a spatially coloured Langevin noise force, with a non-zero spatial range of correlations, since internal bending friction is fundamentally non-local as the force acts between adjacent segments on the chain. In practice, to evaluate the correlation function $\gamma_B(s-s')$, to avoid an ultraviolet catastrophe, we would need to cut the sum off at some sensible upper mode number, corresponding to the smallest relevant length scale, which would cause bending dissipation, such as the base-pair spacing.

### End-to-End friction of a WLC with bending friction

In this section we will calculate the end-to-end friction of a WLC with bending friction by calculating the end-to-end correlation function and relate the friction to the initial slope of the correlation function. The end-to-end friction can also be calculated from the velocity autocorrelation function, using the Green-Kubo integral relation to calculate the effective end-to-end friction. Both give the same answer.

*Calculating the autocorrelation function of the end-to-end distance $\Delta R(t)$*

Firstly, the end-to-end distance of the chain is $R(t) = L - \Delta R(t)$, where at high stretch

$$\Delta R(t) = \frac{1}{2}\int_0^L \mathrm{d}s|\boldsymbol{u}_\perp(s,t)|^2 \; = \; \frac{1}{2}\int_0^L \left(\theta^2(s,t) + \phi^2(s,t)\right) \tag{S11}$$

$$= \; \frac{L}{2}\sum_{n=0}^{\infty}\left(\theta_n^2(t) + \phi_n^2(t)\right), \tag{S12}$$

where $\boldsymbol{u}_\perp = (\theta(s,t), \phi(s,t))^T$ and $\theta_n(t)$ and $\phi_n(t)$ are the Fourier amplitudes of two orthogonal components of tangent vector. We now want to calculate the correlation function $\rho_{\Delta R}(t) = \langle \Delta R(t)\Delta R(0)\rangle - \langle \Delta R(t)\rangle\langle \Delta R(t)\rangle$, The first term is

$$\langle \Delta R(t)\Delta R(0)\rangle \; = \; \frac{L^2}{4}\sum_{n=0}^{\infty}\sum_{m=0}^{\infty}\langle\left(\theta_n^2(t) + \phi_n^2(t)\right)\left(\theta_m^2(0) + \phi_m^2(0)\right)\rangle \tag{S13}$$

$$= \; \frac{L^2}{4}\sum_{n=0}^{\infty}\sum_{m=0}^{\infty}\left(\langle\theta_n^2(t)\theta_m^2(0)\rangle + \langle\theta_n^2(t)\phi_m^2(0)\rangle\right. \tag{S14}$$

$$\left. + \langle\phi_n^2(t)\theta_m^2(0)\rangle + \langle\phi_n^2(t)\phi_m^2(0)\rangle\right). \tag{S15}$$

The Fourier mode amplitudes are

$$\theta_n(t) = \frac{q_n}{\zeta_n}\int_{-\infty}^{t}\mathrm{d}t_1 f_n(t_1)e^{-\frac{t-t_1}{\tau_n}} \tag{S16}$$

and

$$\phi_n(t) = \frac{q_n}{\zeta_n}\int_{-\infty}^{t}\mathrm{d}t_1 g_n(t_1)e^{-\frac{t-t_1}{\tau_n}}, \tag{S17}$$

where $\boldsymbol{f}_n(t) = (f_n(t), g_n(t))^T$, with $\langle f_n(t), g_n(t)\rangle = 0$ and we have used the standard Langevin solution to Eqn.S5 and that $\boldsymbol{u}_\perp(s,t) = \partial_s \boldsymbol{R}(s,t)$ to give $(\theta_n, \phi_n)^T =$

$q_n \boldsymbol{r}_n$, where $\boldsymbol{u}_\perp(s,t)$ is expanded as a sine basis. It is clear then that the autocorrelation function couples to fourth order moments of the Langevin noise terms:

$$\langle \theta_n^2(t)\theta_m^2(t')\rangle = \frac{q_n^2 q_m^2}{\zeta_n^2\zeta_m^2}\int_{-\infty}^{t}\int_{-\infty}^{t}\mathrm{d}t_1\mathrm{d}t_2\int_{-\infty}^{t'}\int_{-\infty}^{t'}\mathrm{d}t_3\mathrm{d}t_4\langle f_n(t_1)f_n(t_2)f_m(t_3)f_m(t_4)\rangle e^{-\frac{2t-t_1-t_2}{\tau_n}}e^{-\frac{2t'-t_3-t_4}{\tau_m}}$$

$$\langle \theta_n^2(t)\phi_m^2(t')\rangle = \frac{q_n^2 q_m^2}{\zeta_n^2\zeta_m^2}\int_{-\infty}^{t}\int_{-\infty}^{t}\mathrm{d}t_1\mathrm{d}t_2\int_{-\infty}^{t'}\int_{-\infty}^{t'}\mathrm{d}t_3\mathrm{d}t_4\langle f_n(t_1)f_n(t_2)g_m(t_3)g_m(t_4)\rangle e^{-\frac{2t-t_1-t_2}{\tau_n}}e^{-\frac{2t'-t_3-t_4}{\tau_m}}$$

$$\langle \phi_n^2(t)\theta_m^2(t')\rangle = \frac{q_n^2 q_m^2}{\zeta_n^2\zeta_m^2}\int_{-\infty}^{t}\int_{-\infty}^{t}\mathrm{d}t_1\mathrm{d}t_2\int_{-\infty}^{t'}\int_{-\infty}^{t'}\mathrm{d}t_3\mathrm{d}t_4\langle g_n(t_1)g_n(t_2)f_m(t_3)f_m(t_4)\rangle e^{-\frac{2t-t_1-t_2}{\tau_n}}e^{-\frac{2t'-t_3-t_4}{\tau_m}}$$

$$\langle \phi_n^2(t)\phi_m^2(t')\rangle = \frac{q_n^2 q_m^2}{\zeta_n^2\zeta_m^2}\int_{-\infty}^{t}\int_{-\infty}^{t}\mathrm{d}t_1\mathrm{d}t_2\int_{-\infty}^{t'}\int_{-\infty}^{t'}\mathrm{d}t_3\mathrm{d}t_4\langle g_n(t_1)g_n(t_2)g_m(t_3)g_m(t_4)\rangle e^{-\frac{2t-t_1-t_2}{\tau_n}}e^{-\frac{2t'-t_3-t_4}{\tau_m}}.$$

Using Wick's theorem (cite) for Gaussian distributed random variables $a, b, c, d$:

$$\langle abcd\rangle = \langle ab\rangle\langle cd\rangle + \langle ac\rangle\langle bd\rangle + \langle ad\rangle\langle bc\rangle,$$

each of the fourth order moments can be evaluated through these pair-wise correlations to give:

$$\langle \theta_n^2(t)\theta_m^2(t')\rangle = \langle \phi_n^2(t)\phi_m^2(t')\rangle = (k_BT)^2\left(\frac{q_n^2q_m^2}{\kappa_n\kappa_m} + \frac{2q_n^4}{\kappa_n^2}\delta_{nm}e^{-\frac{2(t-t')}{\tau_n}}\right), \tag{S18}$$

and

$$\langle \theta_n^2(t)\phi_m^2(t')\rangle = \langle \phi_n^2(t)\theta_m^2(t')\rangle = (k_BT)^2\frac{q_n^2q_m^2}{\kappa_n\kappa_m}. \tag{S19}$$

Hence, plugging these into Eqn.S13, we get

$$\langle \Delta R(t)\Delta R(0)\rangle = (Lk_BT)^2 \left( \left( \sum_{n=0}^{\infty} \frac{q_n^2}{\kappa_n} \right)^2 + \sum_{n=0}^{\infty} \frac{q_n^4}{\kappa_n^2} e^{-\frac{2t}{\tau_n}} \right). \tag{S20}$$

The first order moments of the end-end to end vector are easily calculated:

$$\langle \Delta R(t)\rangle = \langle \Delta R(0)\rangle = Lk_BT \sum_{n=0}^{\infty} \frac{q_n^2}{\kappa_n}. \tag{S21}$$

Putting Eqn.S20 and Eqn.S21 together allows calculation of the autocorrelation function:

$$\rho_{\Delta R}(t) = \langle \Delta R(t)\Delta R(0)\rangle - \langle \Delta R(t)\rangle\langle \Delta R(0)\rangle = (Lk_BT)^2 \sum_{n=0}^{\infty} \frac{q_n^4}{\kappa_n^2} e^{-\frac{2t}{\tau_n}}. \tag{S22}$$

We can confirm the validity of this calculation by calculating the elasticity of the end-to-end distance from using the equipartion theorem $\kappa_{\Delta R} = k_BT/\langle\langle \Delta R^2\rangle\rangle$, where the denominator is the variance of the end-to-end distance:

$$\begin{aligned} \langle\langle \Delta R^2\rangle\rangle = \rho_{\Delta R}(t)|_{t=0} &= (Lk_BT)^2 \sum_{n=0}^{\infty} \frac{q_n^4}{\kappa_n^2} && \text{(S23)}\\ &= \frac{(Lk_BT)^2}{L^2} \sum_{n=0}^{\infty} \frac{q_n^4}{(\kappa_B q_n^4 + F q_n^2)^2} && \text{(S24)}\\ &= (k_BT)^2 \sum_{n=0}^{\infty} \frac{1}{(\kappa_B q_n^2 + F)^2}. && \text{(S25)} \end{aligned}$$

In the large chain limit, we can convert the sum to an integral with $q_n \to q$ and mode spacing $\delta q = \pi/L$:

$$\langle\langle \Delta R^2 \rangle\rangle \approx \frac{(k_B T)^2}{\delta q} \int_0^\infty \frac{\mathrm{d}q}{(\kappa_B q_n^2 + F)^2} \tag{S26}$$

$$= \frac{1}{2} \frac{L(k_B T)^2}{\pi \sqrt{\kappa_B} F^{3/2}} \int_{-\infty}^{\infty} \frac{\mathrm{d}x}{(1+x^2)^2} \tag{S27}$$

$$= \frac{L(k_B T)^2}{4\sqrt{\kappa_B} F^{3/2}}, \tag{S28}$$

where the dimensionless integral, with substitution $x = q\sqrt{\kappa_B/F}$ is evaluated as $\pi/2$. Hence,

$$\kappa_{\Delta R} = \frac{k_B T}{\langle\langle \Delta R^2 \rangle\rangle} = \frac{4\sqrt{\kappa_B} F^{3/2}}{L k_B T}, \tag{S29}$$

which agrees with the high stretch limit of the WLC force-extension relation, $\ell(F) = L(1 - \sqrt{k_B T / 4F\ell_p})$, where

$$\kappa_{\Delta R} = \left(\frac{\partial \ell}{\partial F}\right)^{-1} = \frac{4\sqrt{\kappa_B} F^{3/2}}{L k_B T}.$$

*Calculating the end-to-end friction $\zeta_{\Delta R}$*

To calculate the end-to-end friction, $\zeta_{\Delta R}$ we calculate the time-derivative of the autocorrelation function, which is related to the end-to-end friction as follows [5]:

$$\frac{k_B T}{\zeta_{\Delta R}} = \left| \frac{\mathrm{d}}{\mathrm{d}t} \rho_{\Delta R}(t) \right|_{t=0} = 2(L k_B T)^2 \sum_{n=0}^{\infty} \frac{q_n^4}{\kappa_n \zeta_n} \tag{S30}$$

$$= 2(k_B T)^2 \sum_{n=0}^{\infty} \frac{q_n^2}{(\kappa_B q_n^2 + F)(\tilde{\zeta}_s + \zeta_B q_n^4)}. \tag{S31}$$

Again we take the continuum limit with $q_n \to q$ and mode spacing $\delta q = \pi/L$ to convert the summation to an integral:

$$\frac{k_B T}{\zeta_{\Delta R}} \approx \frac{2(k_B T)^2}{\delta q}\int_0^\infty \mathrm{d}q \frac{q^2}{(\kappa_B q^2 + F)(\tilde{\zeta}_s + \zeta_B q^4)} \tag{S32}$$

$$= \frac{2L(k_B T)^2\sqrt{F}}{\pi \kappa_B^{3/2}}\int_0^\infty \mathrm{d}x \frac{x^2}{(1+x^2)(\tilde{\zeta}_s + \frac{\zeta_B F^2}{\kappa_B^2}x^4)} \tag{S33}$$

$$= \frac{L(k_B T)^2\sqrt{F}}{\kappa_B^{3/2}} \frac{\frac{1}{\sqrt{2}}\left(\sqrt{\tilde{\zeta}_s} + \sqrt{\frac{\zeta_B F^2}{\kappa_B^2}}\right) - \sqrt[4]{\frac{\tilde{\zeta}_s \zeta_B F^2}{\kappa_B^2}}}{\sqrt[4]{\frac{\tilde{\zeta}_s \zeta_B F^2}{\kappa_B^2}}\left(\tilde{\zeta}_s + \frac{\zeta_B F^2}{\kappa_B^2}\right)}. \tag{S34}$$

where we have used the change of variable $x = q\sqrt{\kappa_B/F}$ and evaluated the integral in Wolfram Mathematica. Re-arranging this expression, gives the end-to-end friction of the chain:

$$\zeta_{\Delta R}(F) = \frac{\kappa_B^{3/2}}{L(k_B T)^2\sqrt{F}} \frac{\sqrt[4]{\frac{\tilde{\zeta}_s \zeta_B F^2}{\kappa_B^2}}\left(\tilde{\zeta}_s + \frac{\zeta_B F^2}{\kappa_B^2}\right)}{\frac{1}{\sqrt{2}}\left(\sqrt{\tilde{\zeta}_s} + \sqrt{\frac{\zeta_B F^2}{\kappa_B^2}}\right) - \sqrt[4]{\frac{\tilde{\zeta}_s \zeta_B F^2}{\kappa_B^2}}}. \tag{S35}$$

Mathematically, this expression is valid for all forces, even as $F \to 0$, but in practice since the original semiflexible PDE was derived in the high stretch limit, where $F\ell_P \gg k_B T$, this is also the range of validity of this calculation. In this case, we can simplify this expression further by taking this limit, and which gives us our final expression, used to fit the end-to-end friction of DNA in the main text:

$$\zeta_{\Delta R}(F) = \frac{\sqrt{2}\tilde{\zeta}_s^{1/4}\zeta_B^{3/4}}{k_B T L}F. \tag{S36}$$

It is also possible to get the same result using the Green-Kubo theorem that

$$\frac{k_B T}{\zeta_{\Delta R}} = \int_0^\infty \mathrm{d}t \langle \Delta\dot{R}(t)\Delta\dot{R}(0)\rangle,$$

where $\Delta\dot{R}(t)$ is the end-to-end velocity of the chain and which similarly involves using Wick's theorem to calculate fourth order moments of the velocities of the tangent vector mode-amplitudes $\dot{\theta}_n$ and $\dot{\phi}_n$.

Note that this calculation ignores long range hydrodynamic interactions, which would give rise to Stoke's friction on the whole polymer scale, but is nonetheless reasonable when studying the internal dynamics of the polymer when highly stretched. It is for this reason that in the main text we treat the friction here as the excess friction due to bending dissipation ($\Delta\zeta_{DNA}(F) = \zeta_{\Delta R}(F)$ and that we calculate from the data the equivalent quantity which effectively removes the contribution of DNA solvent friction (as well as the background bead friction), which will have a very weak and flat force dependence at high stretch ($F > 0.08$pN).

### S3. PERSISTENCE LENGTH AT NORMAL AND LOW IONIC STRENGTH CONSISTENT WITH PREVIOUS WORK

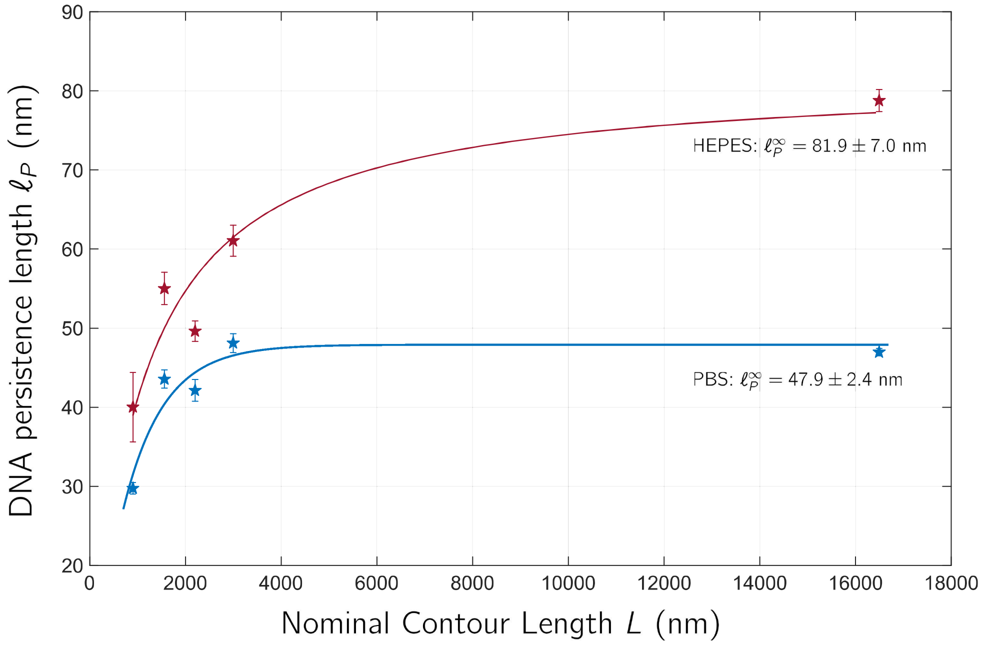


FIG. S12. Plot of mean persistence length measured from force extension experiments as a function of contour length. Error bars are s.e.m from persistence length values from a fit to the extensible WLC model for each single molecule. The number of single molecules used at each length and buffer condition are the same as the main experiment and given in Table.II in the Methods.

For each molecule for which a 5s time series was gathered, we also measured the force extension curve. In addition, we measured the force extension curves for a number of 48kbp DNA, which were not used to calculate the friction as in the main text. In Fig.S12 we plot the mean persistence length of the single molecules of DNA corresponding to different lengths (shown on the $x$-axis) and at two different ionic strengths (using PBS and HEPES buffers). The persistence length is measured by fitting the force-extension curves between 1pN and 10pN

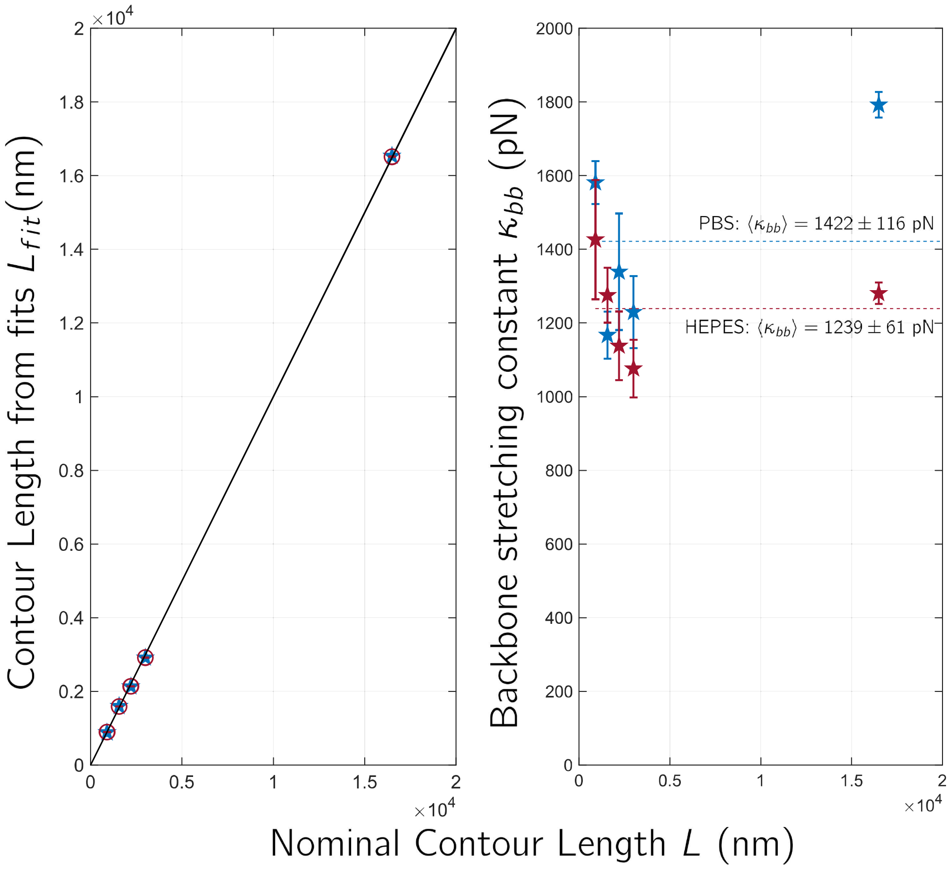


FIG. S13. Plot of mean contour length and stretching elasticity measured from force extension experiments as a function of contour length. Error bars are s.e.m from persistence length values from a fit to the extensible WLC model for each single molecule. The number of single molecules used at each length and buffer condition are the same as the main experiment and given in Table.II in the Methods.

to an extensible WLC equation in the high stretch regime with a force offset $F_0$, where $\ell(F) = L\left(\frac{F'}{\kappa_{bb}} + 1 - \sqrt{\frac{k_B T}{4F'\ell_P}}\right)$ [15], where $F' = F - F_0$. We see that the persistence length increases with increasing contour length, which has been previously observed [20], which is due to finite size effects, such as the role of bead fluctuations. Here we find the PBS data fits an exponentially saturating function $\ell_P(L) = \ell_P^\infty(1 - e^{L/L^*})$, whilst the HEPES data fit a Hill-type saturating function

$\ell_P(L) = \frac{\ell_P^\infty}{1+L^*/L}$, where in both cases $\ell_P^\infty$ is the infinite length ("bulk") persistence length and $L^*$ is the length scale over which it reaches this limit. The values obtained from fits for the infinite length persistence length are $\ell_P^\infty = 47.9 \pm 2.4$ nm for PBS and $\ell_P^\infty = 81.9 \pm 7.0$ nm for HEPES (s.e.m), which are consistent with previous measurements showing an increase in persistence length with decreasing salt/ionic strength [2, 3, 6, 21].

The quantitative values of the contour length and stretching elasticities are shown in Fig.S13, which show that 1) the fitted contour lengths follow closely the nominal lengths, and 2) the stretching elasticities of the backbone are consistent with previous measurements [2, 21].

### S4. RELATIVE IMPORTANCE OF BENDING VS SOLVENT FRICTION FROM MODE ANALYSIS AND RELATION TO A FRICTIONAL FREELY JOINTED CHAIN

A linear force-dependence on friction also arises for a Frictional Freely Jointed Chain (FFJC) [9], which has an angular friction $\zeta_\phi$ restricting the dynamics of relative bond angles: $\zeta_{FFJC} = \frac{\zeta_\phi}{2k_B T N b} F$, where $N$ is the number of links in the FJC. This can be understood intuitively, since contour lengths much greater than the persistence length the DNA chain is essentially freely jointed. It is clear we can make the correspondence that $Nb \leftrightarrow L$, and $\zeta_\phi \leftrightarrow \sqrt{2}\tilde{\zeta}_s^{1/4}\zeta_B^{3/4}$, where the former has a trivial interpretation. The latter quantity arises as the effective asymptotic friction of a dissipative WLC, when solvent friction dominates the slowest mode.

To show this, we note that from Eqn.1 the mode analysis (Eqn.S6&S7) give rise to two lengths scales:

1) $\lambda_\kappa$, below which chain fluctuations are dominated by bending elasticity on length scales less than $\lambda_\kappa$ and by tension $F$ for larger length scales, where from Eqn.S7

$$\lambda_\kappa = 2\pi\sqrt{\frac{\kappa_B}{F}}.$$

So on longer length scales than $\lambda_\kappa$ the WLC fluctuations are not strongly effected by local chain bending and is effectively freely jointed.

2) $\lambda_\zeta$, below which bending friction dominates and above which solvent friction dominates, where from Eqn.S6:

$$\lambda_\zeta = 2\pi\sqrt[4]{\frac{\zeta_B}{\tilde{\zeta}_s}}.$$

In principle, combining these there are in total 4 different viscoelastic regimes.

Here we will only consider those relevant to the optical tweezer experiments, with $1 \leq F \leq 5$pN and lengths of DNA used ($900 \leq L \leq 3000$nm): in this case with $\kappa_B = k_B T \ell_P \approx 199$pNnm$^2$ (for $\ell_P = 48$nm), $\zeta_B = 241\mu$g nm$^3$/ms and $\tilde{\zeta}_s \approx 1.2 \times 10^{-6}\mu$g/ms (see below), we have

$$40\text{nm} \leq \lambda_\kappa \leq 89\text{nm},$$

and

$$\lambda_\zeta \approx 750\text{nm},$$

which shows that the elastic length scale is always much less than the frictional.

To arrive at our approximation $\tilde{\zeta}_s \approx 1.2 \times 10^{-6}\mu$g/ms, we first calculated an initial guess for the frictional length scale by $\lambda_{0\zeta} = 2\pi \sqrt[4]{\frac{\zeta_B}{2\pi\eta}} \approx 494$nm, and then calculating $\tilde{\zeta}_s = 2\pi\eta / \ln(\lambda_{0\zeta}/w) \approx 1.2 \times 10^{-6}\mu$g/ms, where the width of DNA is $w \approx 2.4$nm; with this value we arrive at the value $\lambda_\zeta \approx 750$nm above. In actuality, the critical frictional length is a function of contour length, since $\tilde{\zeta}_s = 2\pi\eta / \ln(L/w)$, depends on the contour length of the WLC, since shorter lengths have an increased effective solvent friction per unit length, since at any given point on the chain there is in total fewer hydrodynamic interactions from other segments on the chain, and conversely the effective solvent friction per unit length is smaller for longer contour lengths, since in total there are more hydrodynamic interactions, inducing on average greater correlated motion with the rest of the chain. This means for contour lengths between $L = 10$nm and $L = 10,000$nm, the critical frictional length varies between $\lambda_\zeta = 540$nm and $\lambda_\zeta = 840$nm; however, this variation is small and it is clear that this does not affect our conclusion that $\lambda_\zeta \approx 750$nm.

So for chain lengths $L \gg \lambda_\kappa$, as is the case for $L > 900$nm in the experiments, then the chain is effectively freely jointed, for separations along the backbone $> \lambda_\kappa$. What is the effective friction of these freely jointed segments — i.e. at this length scale and below? In the absence of bending friction terms, Eqn.1 (Methods) is a hyperdiffusion equation, where perturbations propagate along the chain like $s^4 \sim \frac{\kappa_B}{\tilde{\zeta}_s} t$. Hence, perturbations at frequency $\omega = 1/t$ can only penetrate a length

$$s_s^* \sim \sqrt[4]{\frac{\kappa_B}{\tilde{\zeta}_s \omega}},$$

where the subscript $s$ refer to solvent. The effective frequency dependent compliance at high frequencies is then $J(\omega) \sim s_s^*(\omega)/\kappa_B \sim \frac{1}{\kappa^{3/4}\tilde{\zeta}_s^{1/4}\omega^{1/4}}$.

On the other hand if we exclude solvent friction and only consider bending friction —i.e. if $\lambda_\zeta \to \infty$ — then there is only a single timescale for all modes $\tau_n = \zeta_B/\kappa_B$ and the different segments of the chain respond independently to perturbations, so that the proportion of the chain that responds in time $t$, $s/L = 1 - e^{-t/\tau_B}$. So a perturbation at frequency $\omega = 1/t$ gives rise to a length of chain that responds:

$$s_B^* = L(1 - e^{-\frac{1}{\omega\tau_B}}) \approx \frac{L}{\omega\tau_B},$$

where we can take the latter step when $\omega\tau_B \gg 1$. This then gives an effective high frequency compliance $J(\omega) \sim s_B^*(\omega)/\kappa_B \sim \frac{L}{\zeta_B \omega}$, which is the high frequency compliance of a spring and dashpot with effective friction $\zeta_B$.

Now if we have both solvent and bending friction terms in Eqn.1, where $L > \lambda_\zeta$, such that solvent friction dominates the slowest modes, but at frequencies $\omega > \frac{1}{\tau_B}$, the chain will be dominated by bending friction, this corresponds to a maximum accessible length, or penetration depth of:

$$s_s^*\left(\omega = \frac{1}{\tau_B}\right) \sim \sqrt[4]{\frac{\zeta_B}{\tilde{\zeta}_s}} \sim \lambda_\zeta.$$

Now beyond this frequency, bending friction will dominate and the fraction of the chain that can respond will be $\frac{1}{\omega\tau_B}$, but limited to a total length $\sim \lambda_\zeta$. Hence, the length of chain that responds is

$$s_B^* \sim \frac{\lambda_\zeta}{\omega \tau_B} \sim \frac{\kappa_B}{\tilde{\zeta}_s^{1/4} \zeta_B^{3/4}} \frac{1}{\omega}.$$

And so the high frequency dynamic compliance is:

$$J(\omega) \sim \frac{s_B^*}{\kappa_B} \sim \frac{1}{\tilde{\zeta}_s^{1/4} \zeta_B^{3/4}} \frac{1}{\omega},$$

which is the high frequency response of a spring and dashpot with effective friction $\tilde{\zeta}_s^{1/4} \zeta_B^{3/4}$.

Putting this analysis together, for dissipative WLC of contour length $L >$ 900nm — as in our experiments — stretched to forces $F >$ 1pN, then bending elasticity is weak on length scales $> \lambda_\kappa \approx$ 89nm, so the chain is effectively freely jointed on long length scales, since $L \gg \lambda_\kappa$. According to [9] this gives a friction $\zeta_{FFJC}(F) \sim \zeta_\phi F$. The effective friction between these freely jointed segments on the scale of 89nm (or less at higher forces) is dominated by internal friction, as $\lambda_\kappa < \lambda_\zeta$, but mediated by the solvent friction, which imposes a maximum accessible chain length $\sim \lambda_\zeta$ that can respond giving an effective friction $\zeta_\phi \sim \tilde{\zeta}_s^{1/4} \zeta_B^{3/4}$, which agrees with the more exact calculation in the previous section.

### S5. MEAN FIRST PASSAGE TIME (MFPT) FOR LOOP CLOSURE CALCULATIONS

In this section we calculate the MFPT for loop closure under two conditions: 1) where the tangent vectors at segment ends must align (circular loop) and 2) where there is a finite non-zero angle between the tangent vectors $\theta$, which gives a tear-drop conformation for the closed loop. For circular loop closure, we show that when bending elasticity and bending friction dominate, the stochastic dynamics for curvature $\Gamma$ is a simple spring and dashpot, which allows calculation of a simple closed-form solution for the MFPT, whereas previous calculations with semiflexible polymer dynamics that ignore bending dissipation, have very complicated solutions. In the latter case, as has been previously calculated [24], we find the minimum energy conformation corresponds to tear-drop loops with $\theta \approx 100°$, given this we calculate an approximation for the MFPT, which assumes the dominant part of the time is due to areas of the loop with maximum curvature.

Together the circular loop closure and tear-drop loop closure MFPT calculations form lower and upper bounds on empirical loop closure experiments, and as we show in the main text, we find for $\ell_p >= 50$nm our predictions agree very well with these experiments, compared to MPFT calculation based on solvent friction [8, 14], which massively under predict the time.

#### Simple closed form solution for MFPT of circular loop closure when bending friction and elasticity dominate

If we consider the loop-closure dynamics of small segments of DNA, which have contour length of order the persistence length ($L \sim \ell_P$), then we can make the approximation that bending energy dominates chain entropy and that bending friction dominates solvent friction, the conformation of DNA will, approximately, always be an arc of a circle; as the chain fluctuates due to Brownian motion there then is only a single degree of freedom ,the radius of the circular bend $R$, or its curvature $\Gamma = 1/R$, where $R$ is the radius of this circle. The resulting stochastic dynamics is then the standard Smoluchowski equation for a single degree of freedom, the curvature $\Gamma$:

$$\frac{\partial p(\Gamma,t)}{\partial t} = \frac{1}{\zeta_\Gamma}\frac{\partial}{\partial \Gamma}\left(k_B T\frac{\partial p(\Gamma,t)}{\partial \Gamma} + p(\Gamma,t)\frac{\partial U(\Gamma)}{\partial \Gamma}\right) \tag{S1}$$

where $\zeta_\gamma = \zeta_\Gamma = L\zeta_B$ is the effective friction and $U(\Gamma) = \frac{L}{2}\kappa_B\Gamma^2$ is the bending energy of the chain.

If the population of diffusers is well separated from the barrier, then we can use standard flux-over population method and Kramer's approximations to find the mean first passage time (MFPT) $\tau(R^*)$ to reach the critical curvature $\Gamma^* = 1/R^*$. The equilibrium Boltzmann distribution of curvatures is simply Gaussian $p(\Gamma) = \frac{1}{Z}e^{-\beta U(\Gamma)} = \frac{1}{Z}e^{-\frac{1}{2}\beta\kappa_B L\Gamma^2}$, where $\beta = 1/k_BT$ and $Z = \frac{2\pi}{L\ell_P}$ is the partition function, and so the characteristic width of the distribution about the mean $\langle\Gamma\rangle = 0$ is $\delta\Gamma = \frac{1}{\sqrt{L\ell_P}}$. Therefore the condition for the validity of the Kramers' approximation to the MFPT is that $\delta\Gamma \ll \Gamma^*$, or that $R^* \ll \sqrt{L\ell_P}$. For cyclisation, $R^* = \frac{L}{2\pi}$ and so we require $L \ll 4\pi^2\ell_P$. For $\ell_P \approx 48$nm, this means $L \ll 1895$nm, which is much larger than $L \approx 400$nm, which is the upper limit of contour lengths for which bending friction dominates.

The Kramers' approximation to the MFPT is then given by

$$\tau = \frac{k_BT}{\zeta_\Gamma}ZZ_b,$$

where $Z$ is the partition function from the Bolzmann distribution and

$$Z_b = \int_0^{\Gamma^*} \mathrm{d}\Gamma e^{\beta U(\Gamma)}$$

is a Boltzmann-like integral over the barrier, which in this case is a cusp potential with a finite derivative $\chi = \left|\frac{\mathrm{d}U}{\mathrm{d}\Gamma}\right|_{\Gamma=\Gamma^*}$ at the barrier, instead of the usual smooth barrier. For a cusp potential it is standard approximate $Z_b$, by calculating a 1st order Taylor series of the potential about the barrier, such that $U(\Gamma) \approx U(\Gamma^*) + \chi \times (\Gamma - \Gamma^*) = \frac{1}{2}L\kappa_B(\Gamma^*)^2 + \chi \times (\Gamma - \Gamma^*)$ to give

$$Z_b = e^{\frac{1}{2}\beta L\kappa_B(\Gamma^*)^2} \int_0^{\Gamma^*} \mathrm{d}\Gamma e^{\beta\chi\times(\Gamma-\Gamma^*)} \approx e^{\frac{1}{2}\beta L\kappa_B(\Gamma^*)^2} \int_{-\infty}^{\Gamma^*} \mathrm{d}\Gamma e^{\beta\chi\times(\Gamma-\Gamma^*)} = \frac{k_B T}{\chi} e^{\frac{1}{2}\beta L\kappa_B(\Gamma^*)^2}.$$

The derivative of the potential at the barrier is $\chi = L\kappa_B\Gamma^*$ and so putting these elements together gives the MFPT to a critical radius $R^*$ as

$$\tau(R^*) = \frac{\zeta_B}{\kappa_B} R^* \sqrt{\frac{2\pi}{L\ell_p}} e^{\frac{L\ell_p}{(R^*)^2}}, \tag{S2}$$

which is only valid in the limits given above. Loop closure corresponds to $\Gamma^* = \frac{2\pi}{L}$, giving a MFPT:

$$\tau(R^* = L/2\pi) = \frac{\zeta_B}{\kappa_B} \sqrt{\frac{L}{2\pi\ell_p}} e^{\frac{2\pi^2\ell_p}{L}}, \tag{S3}$$

#### MFPT for tear-drop conformations of DNA

To calculate the MFPT for DNA when the ends can close with the tangent vector at each end being any angle — corresponding to end-hybridisation experiments this is the most likely occurrence initially, as opposed to the tangent vectors being aligned in a circle, we make the approximation that the most probable initial angle of contact will be the minimum energy conformation. Given the energy of this loop, as a function of bend angle we can then approximate the MFPT,

### Minimum energy tear-drop conformation of WLC

The starting point, is to find the conformation or planar space curve of an elastic rod with bending elastcity $\kappa_B$ such that the ends have a difference in tangent angle $\theta$. This is accomplished by solving the Euler-Lagrange equations for the rod with an effective tension $f$ holding the ends together [18]:

$$\kappa_B \frac{\mathrm{d}^2\phi}{\mathrm{d}s^2} + f\sin\left(\phi(s)\right) = 0 \tag{S4}$$

which is the non-linear pendulum equation and where $\phi(s)$ is the tangent vector of the space-curve, which we assume lies in a 2D plane. This is solved with the boundary condition that

$$\phi(L) - \phi(0) = \theta \tag{S5}$$

and that the space-curve must close or touch at its ends:

$$\int_0^{L/2} \mathrm{d}s \cos\left(\phi(s)\right) = 0, \tag{S6}$$

where by symmetry considerations, we expect the solution to be odd about $L/2$, we only need consider the cosine of the tangent angle up to $L/2$. The solution to Eqn.S4 is known to be a Jacobi amplitude elliptic function:

$$\phi(s) = 2am\left(\frac{k}{\sqrt{m}}\left(s - \frac{L}{2}\right)\right) + \pi \tag{S7}$$

where $k = \sqrt{\frac{f}{\kappa_B}}$ and $m$ are parameters of the function, which must be fixed by the boundary conditions.

The condition of a fixed tangent angle is $\phi(0) = \theta/2$ gives the condition:

$$am\left(\frac{kL}{\sqrt{m}}, m\right) = \frac{\pi}{2} - \frac{\theta}{4}. \tag{S8}$$

The loop closure integral gives:

$$\begin{aligned}
\int_0^{L/2} \mathrm{d}s \cos\left(\phi(s)\right) &= \int_0^{L/2} \mathrm{d}s \cos\left(2am\left(\frac{k}{\sqrt{m}}\left(s-\frac{L}{2}\right)\right)+\pi\right) \\
&= -\int_0^{L/2} \mathrm{d}s \left(1-2sn^2\left(\frac{k}{\sqrt{m}}\left(s-\frac{L}{2}\right)\right)\right) \\
&= \frac{kL}{\sqrt{m}}(m-2)+4\mathcal{E}\left(\frac{kL}{2\sqrt{m}},m\right)=0, \qquad \text{(S9)}
\end{aligned}$$

where $\mathcal{E}(u,m)=E(am(u,m),m)$ is the Jacobi Epsilon function (where $E$ is the incomplete elliptic integral of the second kind) and we have used the identities $dn^2(u,m)+m\ sn^2(u,m)=1$ and $\mathcal{E}(u,m)=\int_0^u \mathrm{d}u' dn^2(u',m)$[4], where $dn(u,m)$ is the delta amplitude Jacobi elliptic function. Equations S8 and S9 are then solved numerically using *fsolve.m* function in Matlab [13] to find $k$ and $m$ for a given $\theta$.

Using the same integral identity and that $\frac{\mathrm{d}\ am(u,m)}{\mathrm{d}u}=dn(u,m)$ it is straightforward to calculate the energy of the loop as

$$U=\frac{1}{2}\kappa_B\int_0^L \mathrm{d}s\left(\frac{\mathrm{d}\phi}{\mathrm{d}s}\right)^2=\frac{4\kappa_B k}{\sqrt{m}}\mathcal{E}\left(\frac{kL}{2\sqrt{m}},m\right). \qquad \text{(S10)}$$

On the other hand the energy of a circular loop is $U_0=\frac{L}{2}\kappa_B(\frac{2\pi}{L})^2=2\pi^2\kappa_B/L$, and therefore

$$\frac{U}{U_0}=\frac{2kL}{\pi^2\sqrt{m}}\mathcal{E}\left(\frac{kL}{2\sqrt{m}},m\right),$$

which only depends on the dimensionless combination $kL/\sqrt{m}$, which controls the scale-independent shape of the rod for a given angle $\theta$. In fact, numerically we find $m^*\approx 1.2173$ independent of $L$ and $k^*\approx 4.6754/L$. We plot $U/U_0$ as function of $\theta$ for different contour lengths in Fig.S14, which shows they all collapse to the same curve and we determine by numerically minimisation (Nelder-Mead simplex method using *fminsearch.m* in Matlab) the minimum energy tear-drop conformation to correspond to $\theta^* = 98.58°$, which is close to $\theta \approx 100°$.

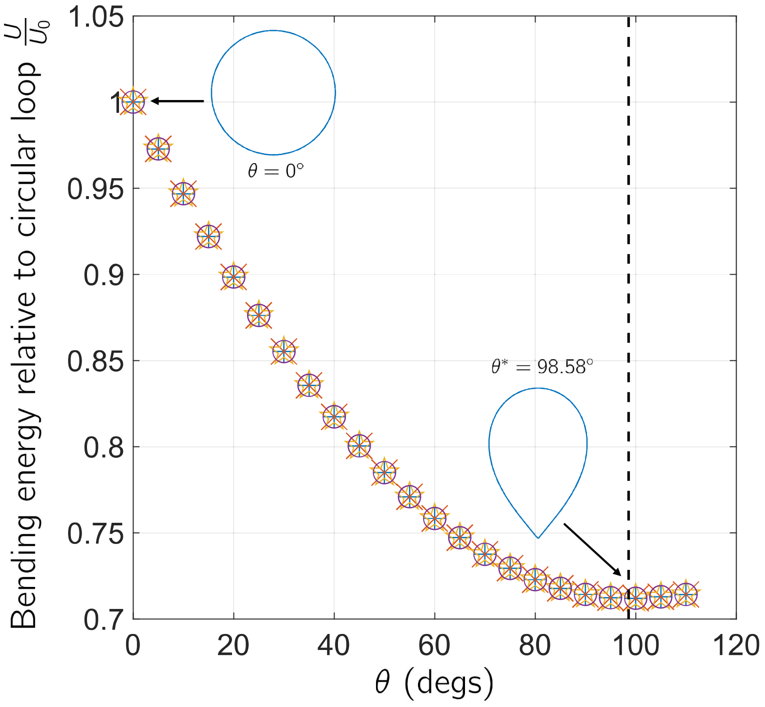


FIG. S14.

#### MFPT to reach looped tear-drop conformation

To approximate the MFPT for loop closure in the tear-drop conformation, we assume that the part of the chain that will dominate the MFPT will be the region of the chain around its centre $s = L/2$. For simplicity, we assume that the threshold is half-maximum curvature; for this central region along the backbone, $L_c$, we calculate the average curvature $\langle \Gamma \rangle_c$. To calculate the MFPT, we then set the critical curvature $\Gamma^* = \langle \Gamma \rangle_c$ and the contour length to $L_c$. This is shown in Fig.S15 for an example of WLC with contour length $L = 100$nm, where for the

455 minimum energy tear-drop conformation ($\theta = \theta^*$), we find $L_c = 53.2$nm and an
456 average curvature along this region corresponding to a circle of radius 14.7nm.

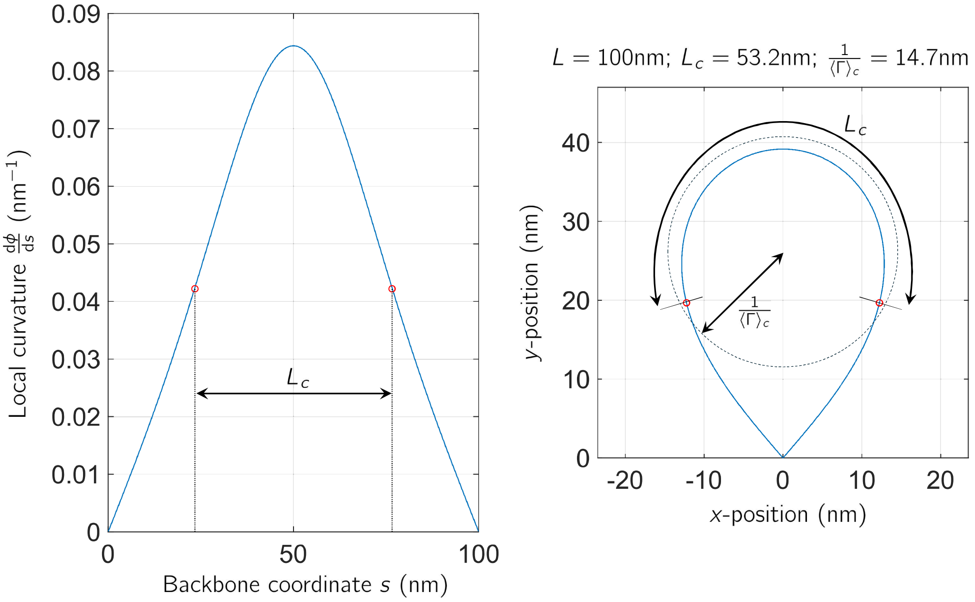


FIG. S15.

### S6. FORCE CORRECTION ON OPTICAL TWEEZERS

At small trap separations, it is known that there are force perturbations arising from a cross-talk between the two traps [12], which we characterise by making measurements of the force on each bead — without DNA — at various trap separations $X$. In Fig.S16, we have plotted 10 replicates of this force over and we see an increasing force for small separations $X$. The forces we measure are qualitatively similar to those measured and calculated by Ling et al [12], where there is exponential decay and oscillatory behaviour for increasing $X$, and the magnitude of forces are of a similar order of magnitude. We fit the average over these 10 replicates in Fig.S16 using the following function, which matches the underlying trend of the data very well, as shown by the solid black line:

$$\delta F(X) = e^{-\frac{X}{\lambda}} \left( f_1 \cos\left(\frac{2\pi X}{\ell}\right) + f_2 \right) + f_0; \tag{S1}$$

with the values of the parameters as given in Table.S1. We see that at small separations $X$ that the force is positive, here defined to be such that the beads are effectively attracted to each other. We also see that the correction should only be significant for 2.6kbp lengths of DNA ($L \approx$ 900nm$\rightarrow$ $\delta F \approx$ 0.5pN) and for the longer lengths (4.5kbps and above, or $L >$ 1500nm) the correction will be $\delta F <$ 0.1pN. This functional form with these fit parameters is then used to correct the forces measured by bead deflections for arbitrary forces.

| Parameter | Value from fits to data in Fig.S16 |
|---|---|
| $\ell$ | 537.69 nm |
| $\lambda$ | 476.19 nm |
| $f_1$ | 0.5036 pN |
| $f_2$ | 3.3095 pN |
| $f_0$ | -0.0383 pN |

TABLE S1. Parameters of Eqn.S1 of fit to data in Fig.S16 of forces between free beads as function of trap separation $X$

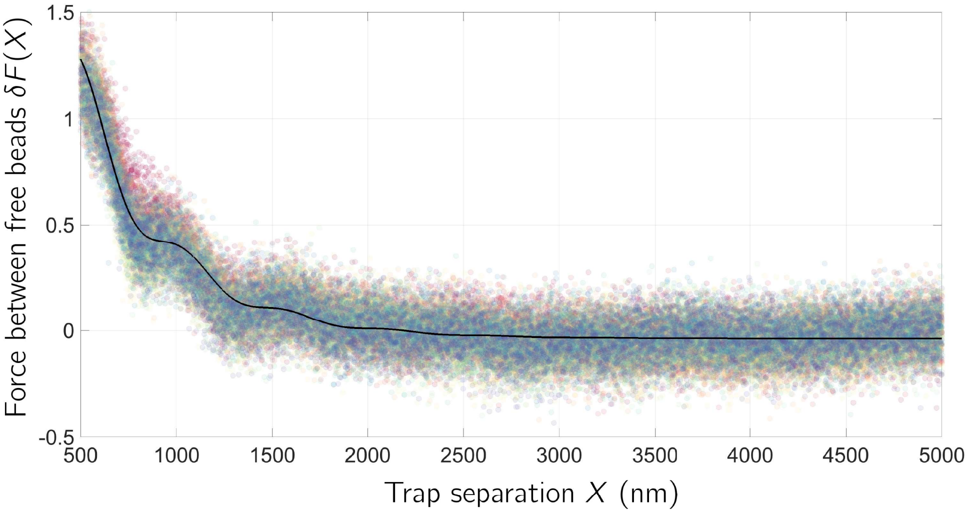


FIG. S16.

[1] M. Atakhorrami, D. Mizuno, G. H. Koenderink, T. B. Liverpool, F. C. MacKintosh, and C. F. Schmidt. Short-time inertial response of viscoelastic fluids measured with Brownian motion and with active probes. *Physical Review E*, 77(6):061508, 2008.

[2] Christoph G. Baumann, Steven B. Smith, Victor A. Bloomfield, and Carlos Bustamante. Ionic effects on the elasticity of single DNA molecules. *Proceedings of the National Academy of Sciences*, 94(12):6185–6190, 1997.

[3] Huimin Chen, Steve P. Meisburger, Suzette A. Pabit, Julie L. Sutton, Watt W. Webb, and Lois Pollack. Ionic strength-dependent persistence lengths of single-stranded RNA and DNA. *Proceedings of the National Academy of Sciences*, 109(3):799–804, 2012.

[4] *NIST Digital Library of Mathematical Functions*. `https://dlmf.nist.gov/22.16.E17`, Release 1.2.4 of 2025-03-15. F. W. J. Olver, A. B. Olde Daalhuis, D. W. Lozier, B. I. Schneider, R. F. Boisvert, C. W. Clark, B. R. Miller, B. V. Saunders,

H. S. Cohl, and M. A. McClain, eds.

[11] L.D.Landau and E.M. Lifshitz. *Fluid Mechanics.* Pergamon Press, 2 edition, 1987.

[12] Lin Ling, Fei Zhou, Lu Huang, Honglian Guo, Zhaolin Li, and Zhi-Yuan Li. Perturbation between two traps in dual-trap optical tweezers. *Journal of Applied Physics*, 109(8):083116, 2011.

[13] MATLAB. *24.1.0.2603908 (R2024a).* The MathWorks Inc., Natick, Massachusetts, 2024.
